# The coiled-coil domain of *E. coli* FtsLB is a structurally detuned element critical for modulating its activation in bacterial cell division

**DOI:** 10.1101/2021.04.21.440662

**Authors:** Samuel J. Craven, Samson G.F. Condon, Gladys Diaz-Vazquez, Qiang Cui, Alessandro Senes

## Abstract

The FtsLB complex is a key regulator of bacterial cell division, existing in either an *off* or *on* state which supports the activation of septal peptidoglycan synthesis. In *Escherichia coli*, residues known to be critical for this activation are located in a region near the C-terminal end of the periplasmic coiled-coil domain of FtsLB, raising questions about the precise role of this conserved domain in the activation mechanism. Here, we investigate an unusual cluster of polar amino acids found within the core of the FtsLB coiled coil. We hypothesized that these amino acids likely reduce the structural stability of the domain and thus may be important for governing conformational changes. We found that mutating these positions to hydrophobic residues increased the thermal stability of FtsLB but caused cell division defects, suggesting that the coiled-coil domain is a “detuned” structural element. In addition, we identified suppressor mutations within the polar cluster, indicating that the precise identity of the polar amino acids is important for fine-tuning the structural balance between the *off* and *on* states. We propose a revised structural model of the tetrameric FtsLB (named the “Y-model”) in which the periplasmic domain splits into a pair of coiled-coil branches. In this configuration, the hydrophilic terminal moieties of the polar amino acids remain more favorably exposed to water than in the original four-helix bundle model (“I-model”). We propose that a shift in this architecture, dependent on its marginal stability, is involved in activating the FtsLB complex and triggering septal cell wall reconstruction.

## Introduction

Cell division in bacteria is a complex process involving intricate coordination between numerous cellular components. Central to this coordination is the divisome, a multiprotein complex that in the gram-negative bacterium *Escherichia coli* consists of a number of essential proteins (FtsZ, FtsA, ZipA, FtsE, FtsX, FtsK, FtsQ, FtsL, FtsB, FtsW, FtsI, and FtsN; Fig. 1) as well as a suite of nonessential, conditionally essential, and redundant proteins^1, 2^. These proteins mediate the various functions necessary for cell division, including establishing the site of division, coordinating invagination of the inner and outer membranes, and remodeling the cell wall at midcell into a septum to compartmentalize the nascent daughter cells. If any of these functions is abrogated through deletion or mutation of essential proteins, the bacteria can continue to elongate and replicate their DNA, but they will be unable to divide and form distinct daughter cells. This will result in the formation of long filaments and eventual cell lysis and death.

**Fig. 1.**
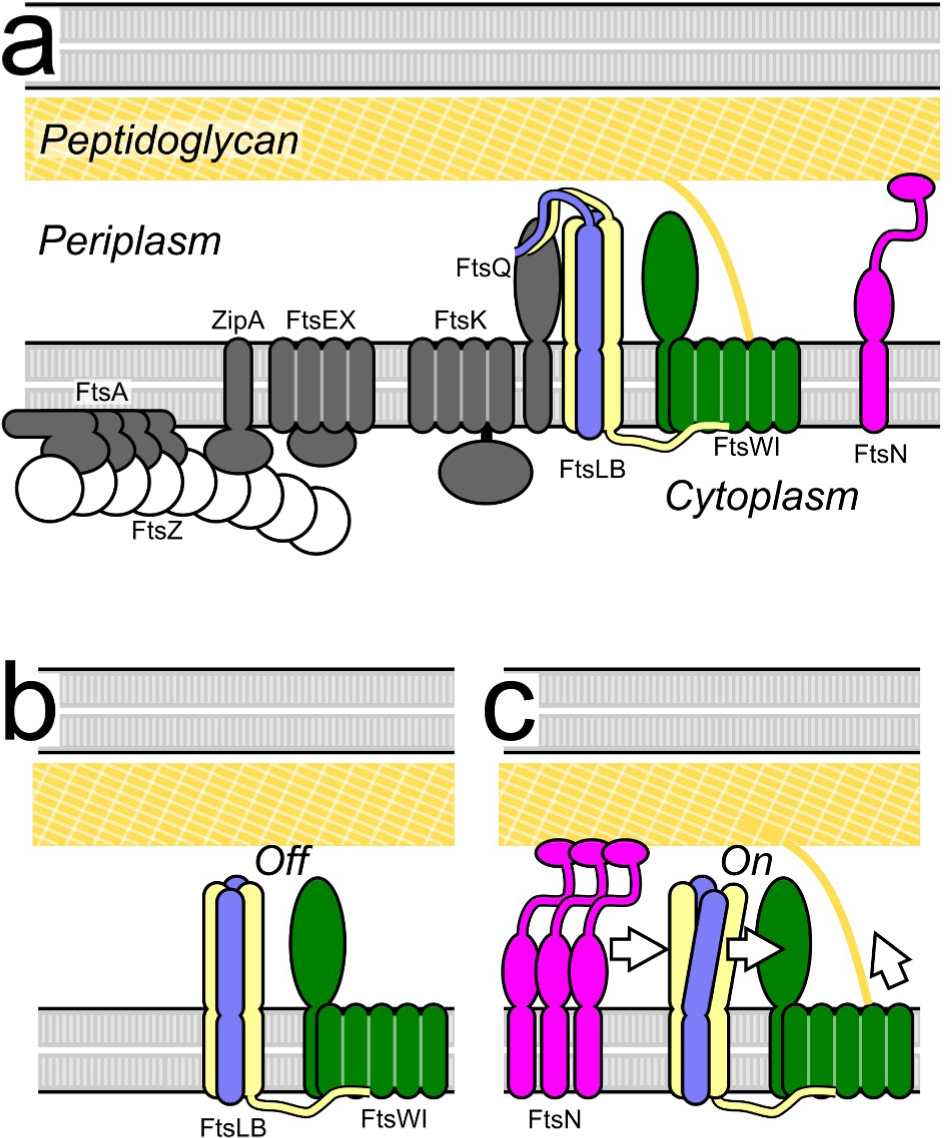
The current model for activation of cell division. a) Schematic representation of the essential components of the *E. coli* divisome. The polymeric ring formed by FtsZ establishes the site of division in coordination with other early components. The peptidoglycan synthase complex FtsWI is the main actor for the reconstruction of the cell wall, leading to the formation of a septum and, eventually, the poles of the nascent daughter cells. Its activation is tightly regulated by FtsN and FtsLB. b) In the current model, FtsWI is in complex with FtsLB but is initially inactive. c) Accumulation of FtsN at midcell somehow triggers a conformational change in the FtsLB complex, which in turn triggers peptidoglycan synthesis likely by a direct interaction with FtsWI.

Remodeling of the cell wall during division involves degradation of old peptidoglycan (PG) at the division site and synthesis of new material, leading to the formation of a septum that eventually splits into the poles of the nascent daughter cells. Numerous nonessential and redundant enzymes (e.g., periplasmic hydrolases) participate in PG reconstruction^3–5^, but the major synthetic activity is performed by the essential FtsWI complex^6–8^. FtsW – a large multipass membrane protein – is a PG glycosyltransferase that polymerizes novel glycan strands from lipid II precursors^9, 10^ (FtsW was also previously reported to have lipid II flippase activity^11–13)^. FtsI is the cognate transpeptidase of FtsW and is responsible for crosslinking the glycan polymers to form a network of PG strands^6, 14^. Forming the septum requires other PG synthases (e.g., either of the bifunctional glycosyltransferase/transpeptidases PBP1a or PBP1b^15^), but functional redundancy between these proteins means that no individual component is essential apart from FtsWI.

The mere presence of FtsWI at midcell is not, however, sufficient for completion of cytokinesis. Instead, the complex must be switched on, and this activation (along with the consequent triggering of cell wall reconstruction) is a tightly regulated step of cell division^1, 16, 17^. In current models, activation begins with the midcell localization of FtsN^18–20^ (Fig. 1c), which communicates with FtsWI through a cytoplasmic route involving FtsA^21–23^ and through a periplasmic route involving the FtsLB complex^21, 24–27^. In this work, we focus on the latter periplasmic activation route.

FtsL and FtsB are both single-pass membrane proteins with a short (FtsL) or minimal (FtsB) N-terminal tail in the cytoplasm and a larger C-terminal domain in the periplasm. They form a heterotetrameric complex consisting of two FtsL and two FtsB subunits arranged into a long helical bundle formed by its transmembrane and periplasmic coiled-coil domains ^28, 29^ (Fig. 2a). The N-terminal, cytoplasmic tail of FtsL is involved with recruiting FtsWI to the division site^30^, whereas the C-terminal, periplasmic tail of FtsB binds with high affinity to another divisome protein FtsQ^31–33^ and is needed for FtsLB’s own recruitment to midcell^30, 34–36^. The structure of FtsLB has not been solved experimentally aside from fragments^31, 32, 37^, but computational structural models of its helical bundle region are available. Originally, the Monasterio group proposed models of the soluble coiled-coil region in complex with the periplasmic domain of FtsQ, in either trimeric or hexameric configurations^38^. More recently, using a set of amino acid contacts postulated by evolutionary coupling, we derived a model that includes both the transmembrane region and the coiled coil^29^. In this model, the transmembrane and coiled-coil domains of FtsL form a continuous helix, whereas FtsB contains a potentially flexible, Gly-rich linker that breaks the helix between these two regions^37^.

**Fig. 2.**
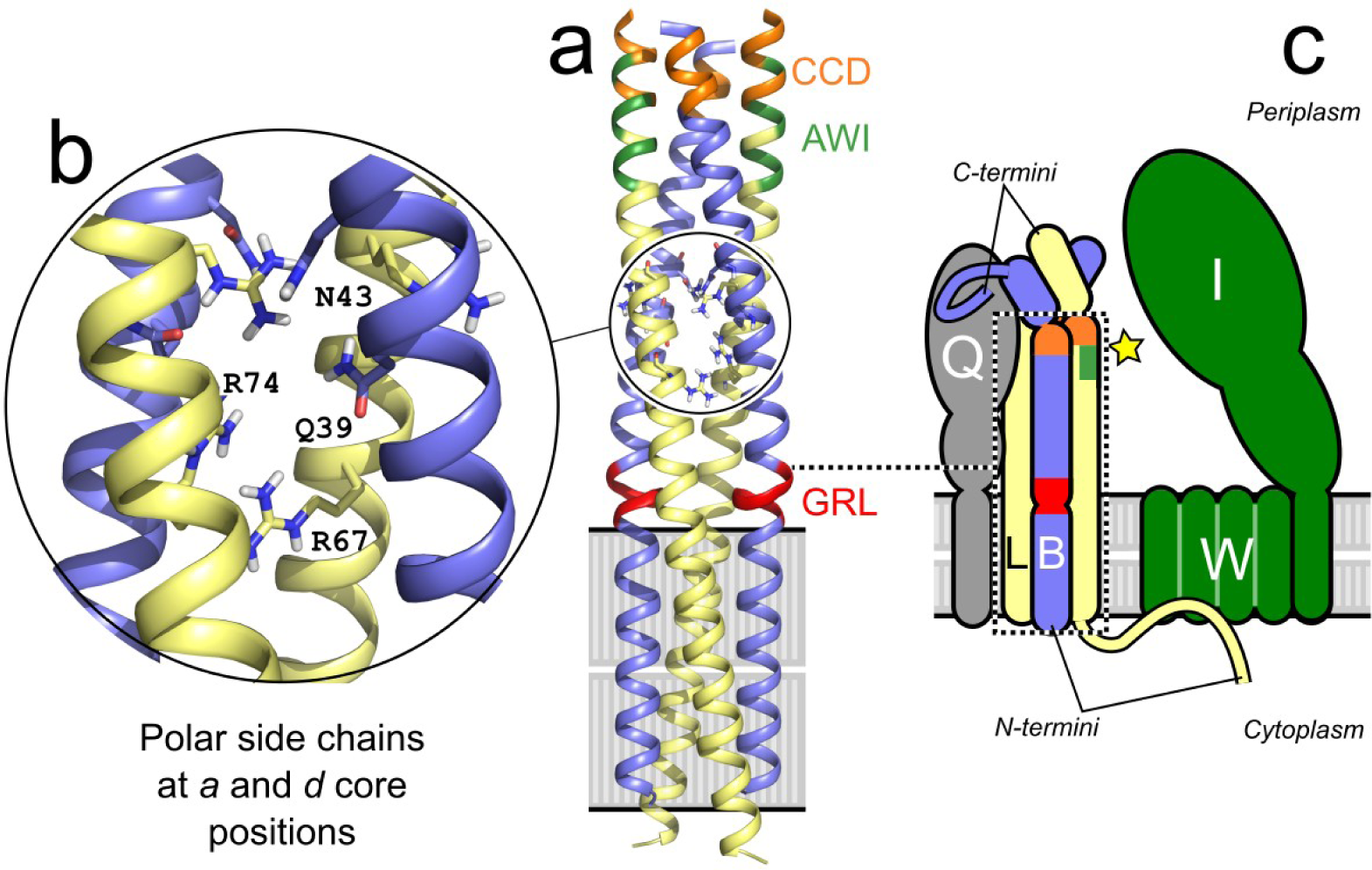
An unusual cluster of strongly polar amino acids in the core of the computational model (I-model) of FtsLB likely tunes its stability and conformation dynamics. a) Original computational model of the tetrameric helical bundle of FtsLB^29^, referred to here as the I-model. The model is formed by two FtsL (yellow) and two FtsB (blue) units. FtsL forms a continuous helix across the membrane and periplasmic domains, whereas the helix of FtsB is interrupted in the juxta-membrane region by an unwound Gly-rich linker (GRL, red). Highlighted in orange and green are the CCD and AWI regions, respectively, which occur near a predicted hinge at the C-terminal end of the coiled coil. These regions are critical for FtsLB activation. b) The coiled coil contains a cluster of polar amino acids at core “*a*” and “*d*” positions, including two Arg residues in FtsL (positions 67 and 74) and Gln-39 and Asn-43 in FtsB. The polar cluster is likely to be a destabilizing feature of the coiled coil. c) Schematic representation of FtsLB and its interactions with FtsQ and the FtsWI complex. The region corresponding to the model is enclosed in the dotted box. The interaction with FtsQ is largely mediated by the C-terminal tail of FtsB. The interaction with FtsW is mediated by the N-terminal tail of FtsL. The star indicates the putative activating contact between the AWI region of FtsL and FtsI.

In current models, FtsLB regulates FtsWI septal PG synthesis activity by transitioning from an *off* state to an *on* state in response to a signal from FtsN (Fig. 1c). This idea was initially proposed following the identification of a series of gain-of-function mutations within both FtsL and FtsB that enable survival in the absence of the normally essential FtsN ^21, 24^. Along with subsequent work^25, 26^, this led to the identification of two related regions at the C-terminal end of the FtsLB coiled coil that are central to its regulation of FtsWI. The first region, named the Constriction Control Domain (CCD, approximately residues 88-94 in FtsL and 55-59 in FtsB), houses the aforementioned *ΔftsN*-suppressing mutations^21, 24^. The second region neighbors the CCD on the opposite helical face of FtsL specifically and is designated as <u>A</u>ctivation of Fts<u>WI</u> (AWI; positions 82-84, 86-87, and 90). Dominant-negative mutations in FtsL indicate that the AWI region directly interacts with and activates FtsWI^25, 26^, suggesting that the *off/on* transition in FtsLB may involve conformational changes that make the AWI region available to interact with and activate FtsWI^26^. Normally, either direct or indirect interactions with FtsN are required to trigger such a change; however, the gain-of-function CCD mutations may induce similar structural rearrangements of FtsLB, thereby mimicking the signal from FtsN and bypassing its requirement to trigger septal PG reconstruction. Whatever conformational changes in FtsLB are required for this activation, they will likely depend on the extended helical topology that is at the core of the complex^29, 35, 39, 40^.

One intriguing structural feature of FtsLB is the unusual presence of a cluster of strongly polar amino acids buried at interfacial positions of the coiled coil (i.e., at positions designated as “*a*” and “*d*” in the “*abcdefg*” heptad repeat). This “polar cluster” (as it will be referred to from now on) consists of two arginine residues (R67 and R74) in FtsL and of a glutamine residue (Q39) and two asparagine residues (N43 and N50) in FtsB (Fig. 2b). Since canonical coiled coils contain primarily hydrophobic residues at the interfacial “*a*” and “*d*” positions^41^, the polar cluster is likely to decrease the stability of the coiled coil^29^. This suggests that these nonideal residues play some critical role in modulating the stability and dynamics of the FtsLB complex, which likely has important functional consequences – a hypothesis that we address in this present article.

We hypothesize that the coiled coil of FtsLB is detuned for stability in order to support the dynamics necessary for structural transitions that occur during the *off* to *on* switch at the heart of FtsWI regulation. Here, we show that hydrophobic mutations introduced in the polar cluster of FtsLB stabilize the coil but lead to cell division defects *in vivo*. We also show that the identity of the interfacial residues is important for the observed division phenotypes, indicating those residues may play a more nuanced role than simply destabilizing the hydrophobic coiled-coil interface and are likely to participate in the balance of forces that regulate the *off*/*on* transition of the complex. Additionally, because the presence of polar residues at “*a*” and “*d*” positions statistically favors the formation of two-stranded coiled coils^42–44^, we investigated an alternative model of FtsLB in which the coiled-coil region splits into a pair of two-stranded coiled-coil domains (the Y-model), as opposed to the monolithic, four-stranded coil that we originally proposed^29^ (the I-model). This revised model fits the available evidence as well as the original I-model, while displaying better behavior, and thus we propose it as the more likely candidate for the structural organization of FtsLB.

## Results and Discussion

### The polar cluster of FtsLB is an unusual feature for coiled-coil structures

Although “*a*” and “*d*” positions of coiled coils tend to be mainly hydrophobic, polar amino acids can also occur there. To determine if the coiled coil of FtsLB is unusually rich in interfacial polar amino acids in comparison with other coiled coils, we performed a structural analysis of the 2,662 crystal structures available in the CC+ database^45^. We found that polar amino acids such as Asp, Glu, His, Arg, Lys, Gln, and Asn occur with a frequency of 17.6% at “*a*” or “*d*” positions, corresponding on average to approximately one polar amino acid every three heptad repeats (supplementary Table S1). Notably, the propensity to accommodate polar amino acids decreases as the number of helices forming the coiled-coil assembly increases. The frequency of polar amino acids is highest at 18.8% in coiled coils formed by two helices, and it decreases to 14.1% and 11.7% for three-stranded and four-stranded coiled coils, respectively. In contrast, 30% of the “*a*” and “*d*” positions in *E. coli* FtsLB are polar, confirming that the level of enrichment is quite high in comparison with the average coiled coil and nearly three times the average frequency for coiled coils that assume a tetrameric configuration, suggesting that the coiled coil of FtsLB is not designed for maximal stability.

### The polar cluster is evolutionarily conserved

To investigate if the polar cluster is a conserved feature of FtsLB, we analyzed a multisequence alignment containing 2900 pairs of FtsB-FtsL sequences from diverse proteobacterial taxa. In *E. coli*, the coiled-coil region is predicted to extend for approximately five heptad repeats, starting after the transmembrane helices and ending approximately at the CCD region^29^. Three polar amino acids are found in FtsB at positions 2*d* (Gln-39, the “*d*” position in the second heptad repeat), *3a* (Asn-43), and *4a* (Asn-50). FtsL contains two arginine residues at positions *2a* and *3a* (Arg-67 and Arg-74, respectively). Additionally, FtsL contributes Glu-88 (5a), one of the critical amino acids in the CCD region^21, 24^ at the margin of the coiled coil.

The pattern of polar/nonpolar amino acids found at “*a*” and “*d*” positions in the alignment is summarized graphically in Fig. 3. We found that this feature is highly conserved, even if the specific sequence is not. As highlighted by the main “paths” in the graph, five of the six polar positions in *E. coli* have a strong tendency to be polar in all species. The only position at which a polar amino acid is not particularly conserved corresponds to Arg-67 (2a) of FtsL.

**Fig. 3.**
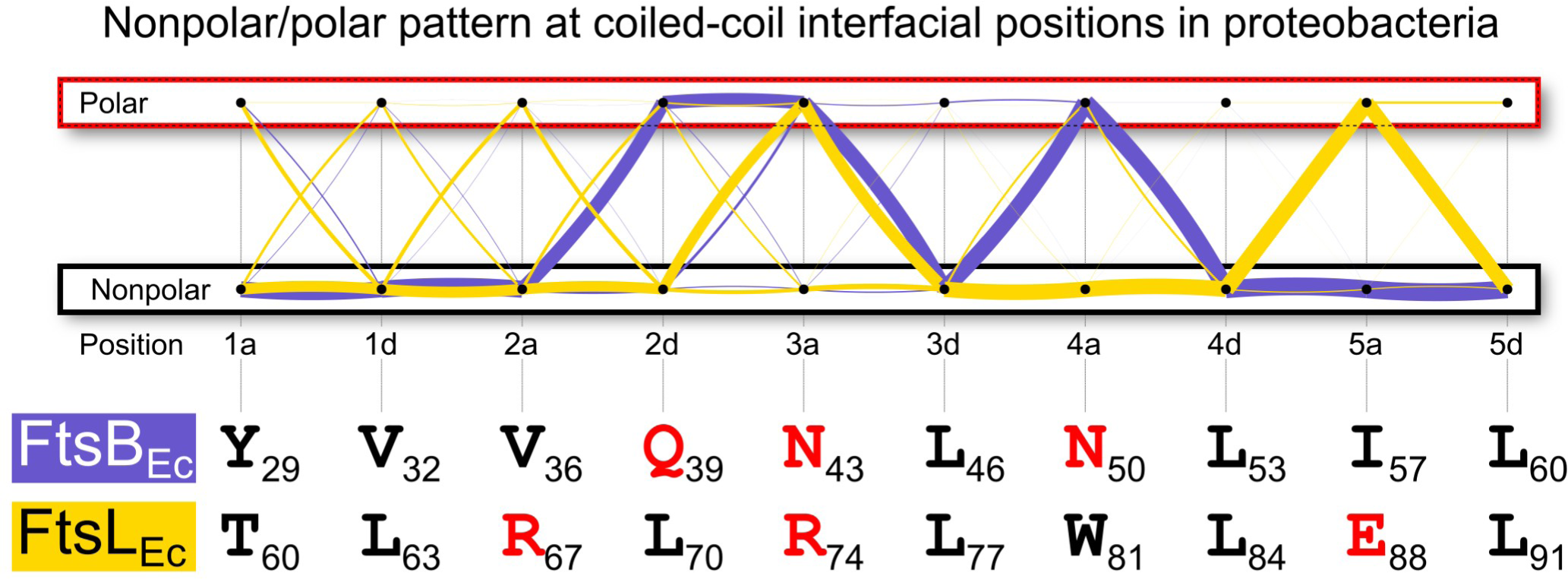
Conservation of the polar cluster in the core “*a*” and “*d*” positions of the coiled coil of FtsLB in proteobacterial species. The yellow (FtsL) and blue (FtsB) lines follow the pattern of polar (Asp, Glu, His, Arg, Lys, Gln, and Asn) and nonpolar amino acids in alignments of 2900 paired FtsB/FtsL **sequences**. The thickness of the lines is proportional to the number of sequences that follows a certain (polar/nonpolar) binary pattern. The graph evidences that polarity at positions corresponding to Gln-39, Asn-43, and Asn-50 is a highly conserved feature in FtsB. Positions Arg-74 and Glu-88 are also most frequently polar in FtsL. Polarity at the position corresponding to Arg-67 is not conserved in FtsL. However, there is overall a 71% probability that at least one position in the first two heptad repeats of FtsL contains another polar residue.

However, a polar amino acid occurs overall frequently across any of the “*a*” and “*d*” positions of the first two heptad repeats (71% of the sequences). In fact, the class of sequences to which *E. coli* belongs (i.e., those sequence in which both FtsL and FtsB contribute three polar amino acids each) is the most common (39%). Moreover, 84% of the sequences contain at least five polar amino acids (supplementary Fig. S1). The analysis therefore suggests that, in proteobacteria, the coiled coil of FtsLB is far from being tuned for maximum stability and is thus designed to be dynamic or metastable for functional reasons.

### The polar cluster tunes FtsLB’s propensity to transition to an activated state

In order to test whether the conserved polar cluster is critical for function, we expressed mutant variants at these positions *in vivo* and examined whether these changes led to cell division defects. In our first round of experiments, we tested the effect of “idealizing” the coiled coil by converting each of the polar amino acids to a canonical hydrophobic residue (Ile or Leu, at “*a*” and “*d*” positions, respectively). We also mutated Trp-81 of FtsL to a small hydrophobic residue, since bulky aromatic residues tend to be excluded from natural coiled-coil interfaces^41^. The effect of individual point mutations was assessed in complementation experiments in which we measured the length distribution of samples of at least 500 cells. As done previously to assess the severity of mutations that cause elongation defects^29^, we measured the proportion of cells with lengths exceeding the 95^th^ percentile of the length distribution observed for the WT (dashed vertical line in the four example length distributions displayed in Fig. 4a-d). The fraction of long cells is reported for further mutants in the histograms in Fig. 4e-h, grouped for FtsB nonideal-to-ideal, FtsL nonideal-to-ideal, FtsL charged-to-charged, and double substitutions at positions R67 and R74 in FtsL. Pictures, cell distributions, and metrics for all mutations are reported in supplementary Fig. S2.

**Fig. 4.**
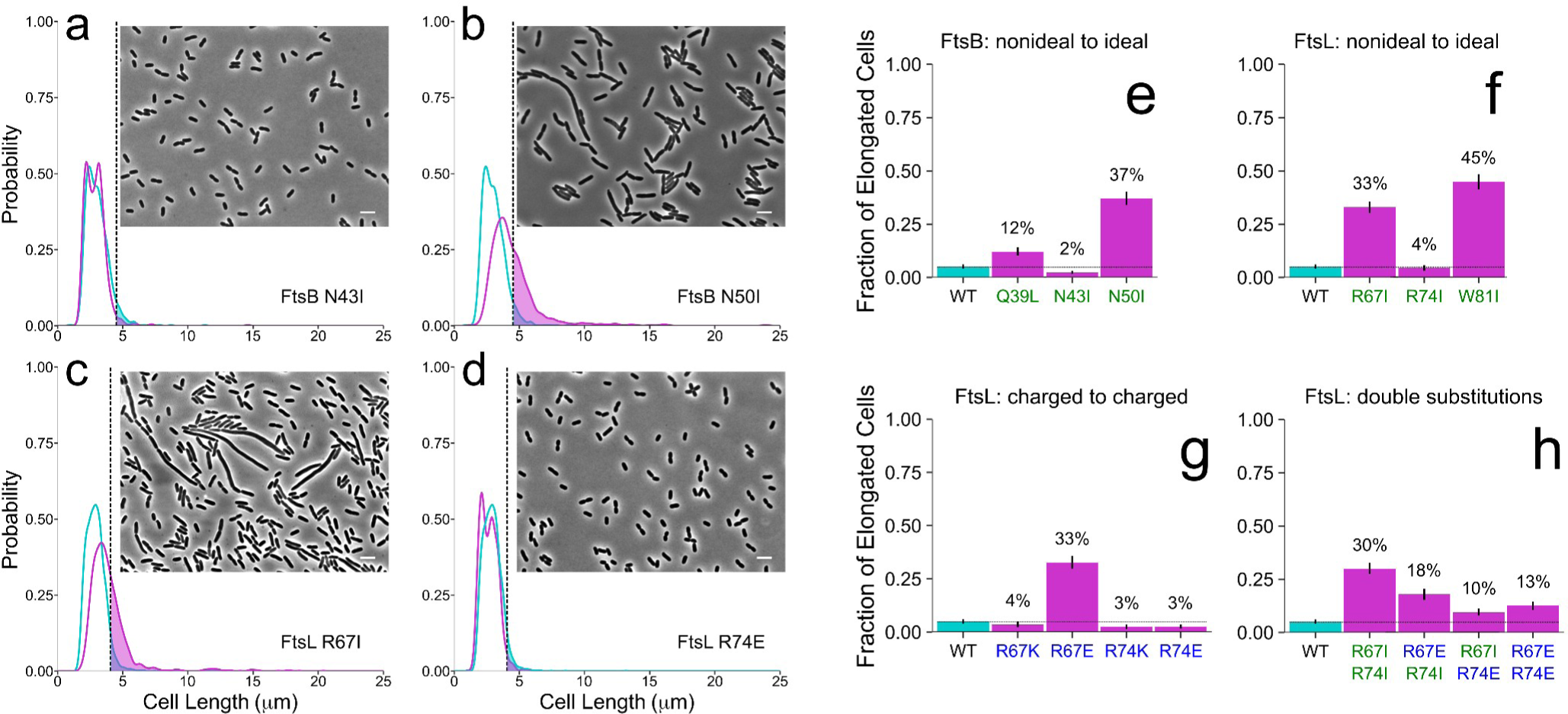
Mutations in the polar cluster cause both elongation and small cell phenotypes *in vivo*. Panels *a*-*d*: Phase-contrast images of representative mutants (5 μm scale bar) and cell length distributions of mutant cells (magenta) compared to wild type (cyan). The shaded areas to the right of the dotted lines represent the elongated cells (i.e., those that are longer than the 95th percentile in the WT distribution). The four examples include (*a*) a wild-type-like mutant (FtsB N43I), (*b-c*) two mutations that cause frequent elongation (FtsB N50I at 37% elongated and FtsL R67I at 33% elongated), and (*d*) a mutant that displays a higher fraction of shorter cells (FtsL R74E). Panels *e*-*h*: histograms of percent of elongated cells for all mutations at the nonideal coiled-coil positions (nonpolar mutations in green; charged mutations in blue). These include (*e*) FtsB and (*f*) FtsL nonideal-to-ideal mutations, (*g*) FtsL charged-to-charged mutations, and (*h*) FtsL double mutations at positions R67 and R74. Nonideal-to-ideal mutations tend to cause notable elongation phenotypes. A remarkable exception is position R74, in which mutations appear to cause a small cell phenotype (likely due to deregulated division). R74E, in particular, suppresses the elongated phenotypes of R67I and R67E, indicating that these positions are important for governing the fine balance between the *on* and *off* conformations of FtsLB. Experiments were performed at 37 °C. Error bars represent the 95% confidence interval of the fraction of elongated cells estimated from 1000 replicates of bootstrap resampling.

Three of the five polar-to-nonpolar individual point mutations tested (FtsB Q39L and N50I; FtsL R67I) resulted in notable defective phenotypes, displaying a fraction between 12-37% of elongated cells (Fig. 4e-f). The remaining two polar-to-nonpolar mutations (FtsL R74I and FtsB N43I) were similar to WT. Mutation of the bulky Trp residue in FtsL (W81I) also resulted in a defective phenotype. To exclude that the division defects were due to changes in protein expression levels, we performed western blot analyses (supplementary Fig. S3), which indicated that each mutant was expressed to similar levels as WT, except for FtsB N43I which showed increased expression. These observations support the hypothesis that the polar cluster plays a critical role in FtsLB function.

Arginine is one of the most polar amino acids, essentially never occurring in its neutral form, and thus it is one of the most destabilizing amino acids when buried within a protein core. For this reason, we further investigated the effects of mutating both Arg-67 and Arg-74 in FtsL (Fig. 4h). The double charged-to-hydrophobic mutation (R67I+R74I) resulted in impaired division in a manner similar to the R67I single mutant (30% vs 33% elongated cells). We then asked whether the identity of these positions is important (Fig. 4g). We first inverted the charge with Glu substitutions. The R67E mutation produced a division-defective phenotype (33% elongated cells) similar to that of R67I. Like the Ile substitution at the same position, R74E did not result in elongation, but somewhat surprisingly, this substitution resulted in a larger fraction of smaller cells (left-shifted peak compared to WT, Fig. 4d). We then preserved the positive charges at the 67 and 74 positions of FtsL with Lys substitutions. While R67K produced WT-like cells, R74K had a similar phenotype to R74E, with a larger fraction of small cells (supplementary Fig. S2c).

The fact that charge reversal mutations at position R67 and R74 result in opposite phenotypes (elongation vs small cells, respectively) is interesting. Gain-of-function mutations in FtsLB that cause early cell division have been identified before^21, 24^, predominantly within the CCD. The reduced cell length of R74 mutants is particularly notable since a similar phenotype was never observed among a total of 55 mutations in the transmembrane region and coiled-coil positions of FtsLB that were assessed in our previous analysis with identical conditions and methodology^29^. For this reason, we combined both charge reversal mutations (R67E+R74E) to observe their interplay. Interestingly, R67E+R74E resulted in a shorter cell length distribution (13% elongated cells compared to 33% for R67E alone, Fig. 4d), supporting the hypothesis that R74E can suppress the R67E elongation defect by inducing early cell division. This is also consistent with the phenotypes we observed in the R67I+R74E and R74E+W81I double mutations (10% and 15% elongated cells, respectively), where the elongated phenotypes of R67I and W81I alone (33% and 45% elongated cells, respectively) are also suppressed, resulting in a decreased fraction of elongated cells.

In order to determine if FtsL R74E is unique or if other mutations at that position can suppress elongation defects, we tested further combinations. Ultimately, we found that the extent of observed suppression varies with the identity of the amino acid at position 74. The charged-to-nonpolar mutation R74I (which was WT-like alone and did not suppress the elongated phenotype of R67I) also appears to reduce the severity of the elongated phenotype when combined with R67E (33% vs 18% elongated cells for R67E and R67E+R74I, respectively). On the other hand, R74K did not noticeably suppress the elongation defect when combined with R67I, R67E, or W81I (supplementary Fig. S2d), suggesting that a negative charge at position 74 (or perhaps merely the lack of a positive charge) is needed for the suppression effect.

Overall, the data suggest that the polar cluster of the FtsLB coiled coil is important for fine-tuning the propensity of the complex to transition to an activated state. In particular, the Arg residues at the 67 and 74 “*a*” positions in FtsL somehow play opposing roles in this regulation, and their specific residue identity is important.

### Hydrophobic substitutions in vitro increase thermal stability

If the polar cluster plays a role in tuning FtsLB’s propensity to transition to an activated state, then polar-to-nonpolar substitutions likely stabilize the coiled-coil domain but may be detrimental for the complex to undergo conformational changes. To directly assess whether the polar cluster does in fact govern the structural stability of FtsLB, we investigated the thermal stability of the complex *in vitro* by monitoring secondary structure content using circular dichroism (CD). In this experiment, we simultaneously mutated all four centrally located polar residues in both FtsB (Q39L and N43I) and FtsL (R67I and R74I).

We used a version of FtsL (FtsL_35-121_) lacking the unstructured N-terminal tail, which is not necessary for the assembly of the FtsLB complex^29^. Both FtsB and FtsL constructs retained the N-terminal purification tags (His and Strep tags, respectively), which do not cleave efficiently. To avoid interference from reducing agents in the sensitive UV region, both native cysteine residues of FtsL were replaced with alanine (C41A and C45A). We have shown previously that this Cys-less construct is stable and functional^29^. The resulting constructs, His-FtsB/Strep-FtsL_35-121_-C41A-C45A and His-FtsB-Q39L-N43I/Strep-FtsL_35-121_-C41A-C45A-R67I-R74I, are termed “WT” and “4×-mutant”, respectively.

Fig. 5a shows the CD spectra of the constructs solubilized in n-dodecyl-β-D-maltopyranoside (DDM) at low temperature (4 °C). Both constructs have the expected spectral signature of helical proteins with high secondary structure content. Little difference is noticeable between the two constructs, suggesting that the four mutations do not cause major changes in helical content at low temperature. We then investigated the difference in stability between WT and 4×-mutant comparing the CD signal at increased temperatures, monitoring ellipticity at one of the helical minima (224 nm, Fig. 5b and supplementary Fig. S4).

**Fig. 5.**
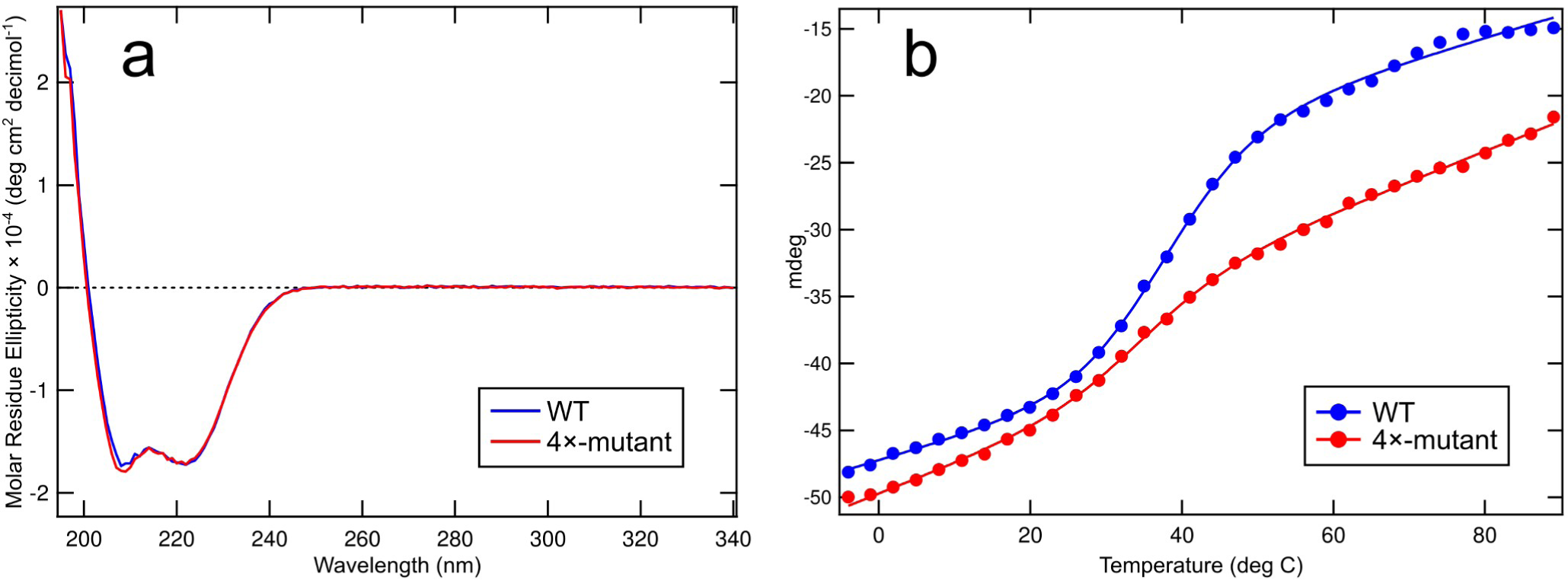
Conversion of the polar cluster to idealized hydrophobic residues increases thermal stability but affects folding cooperativity. a) Far-UV CD spectra of WT FtsLB (blue) compared to the 4×-mutant (red) at 4°C. b) Representative temperature melting curves comparing WT FtsLB (blue) to the 4×-mutant (red). The WT melting curve displays a sigmoidal transition centered around 40 °C. In contrast, the 4×-mutant displays a curve that is shifted to the right but lacks a significant transition, possibly indicating a loss of cooperativity of unfolding. CD scans were monitored at 224 nm from 4 to 89 °C. Replica melting curves are included in supplementary figure Fig. S4.

The WT construct (blue dots) displays a sigmoidal melting curve with a transition centered around 40 °C. Given the high thermal stability of transmembrane helices, the transition is most likely attributable to loss of helicity in the coiled-coil region. The relatively early unfolding transition confirms that the coiled coil of FtsLB is structured but not optimized for stability. The melting curve of the 4×-mutant is markedly different (red dots). The construct retains a higher degree of secondary structure at higher temperatures, indicating that, as expected, the replacement of unfavorable polar side chains for canonical hydrophobic residues stabilized the coiled coil. However, the 4×-mutant’s curve does not display the same degree of cooperativity as the WT, but rather, it shows a nearly linear loss of ellipticity lacking a clear transition. A sharp transition is consistent with a two-state unfolding process typical of well folded proteins; therefore, it is likely that the 4×-mutant version of the coiled coil of FtsLB can access alternative, potentially misfolded conformations. This notion becomes particularly interesting in light of the fact that FtsLB can be modeled in two alternative conformations, as discussed in the next section.

### An alternative structural organization for FtsLB: the Y-model

Experimental evidence *in vitro* indicates that the FtsLB complex is a 2:2 FtsL:FtsB tetramer^28, 29, 37^. We previously modeled the complex in the simplest configuration consistent with this tetrameric state (i.e., a monolithic, four-helix bundle that extended across the transmembrane and periplasmic regions^29^, Fig. 2b). We noted, however, that it is unlikely that a tetrameric coiled coil would be stable with the number of polar residues present in its core. Consistently, the coiled-coil configuration did not appear stable during molecular dynamics simulations, where we observed a marked tendency of this region to open and recruit water within its interior.

The previously discussed analysis of the structural databases indicates that a polar coiled coil would be most likely to assume a two-stranded configuration. Experimental studies on model coiled coils also indicate that polar residues in core positions can influence the number of helical strands in coiled coils, with a two-stranded coiled coil being better suited to accommodate polar residues^42, 43^. In particular, the four Arg residues contributed by the FtsL chains (two from each monomer) are particularly costly to bury, since Arg is essentially always protonated even in a hydrophobic environment. Indeed, a systematic study of all twenty amino acids in a model coiled coil found that Arg is the most destabilizing residue at “*a*” positions and also that, when present, this amino acid strongly favors a two-stranded over a three-stranded configuration^44^. This finding is in good agreement with our analysis of the structural database, which shows that Arg is relatively frequent at “*a*” positions in two-stranded coiled coils (4.5%) but much more rare in four-stranded coils (0.5%, supplementary Table S1). This preference can be explained by the solvent accessibility of the side chains: in two-stranded coiled coils, the narrower interface allows the polar moieties at the end of the side chains to access the surrounding solvent and remain partially water-exposed. As the size of the helical bundle increases, these groups become increasingly more buried, resulting in a higher desolvation cost.

Following the lead suggested by the molecular dynamics analysis of our original FtsLB model^29^, we report a revised model of FtsLB in which the periplasmic region splits into a pair of two-stranded coiled-coil domains, each containing one FtsL and one FtsB chain (Fig. 6a). This model (which we named the “Y-model” from its shape) is based on the same set of side chain contacts between FtsB and FtsL inferred by the co-evolutionary analysis we used to derive the original monolithic model^29^ (named here the “I-model”). The transmembrane region is modeled in the same four-helix-bundle configuration of the original I-model. As in the I-model, the helix of FtsL in the coiled-coil region is continuous with the transmembrane helix, as indicated by experimental analysis^29^. In this configuration, the FtsL helix remains on the outward face of the “Y”, whereas the helices of FtsB occupy the inward face, in close proximity to each other and thus buried within the overall arrangement. With respect to the transmembrane domain, the helix of FtsB is rotated so that its interfacial “ *a*” and “*d*” positions face the corresponding positions of FtsL, a rotation that is readily enabled by the structurally malleable, Gly-rich linker that occurs between the transmembrane and coiled-coil helices of FtsB^29,37^.

**Fig. 6.**
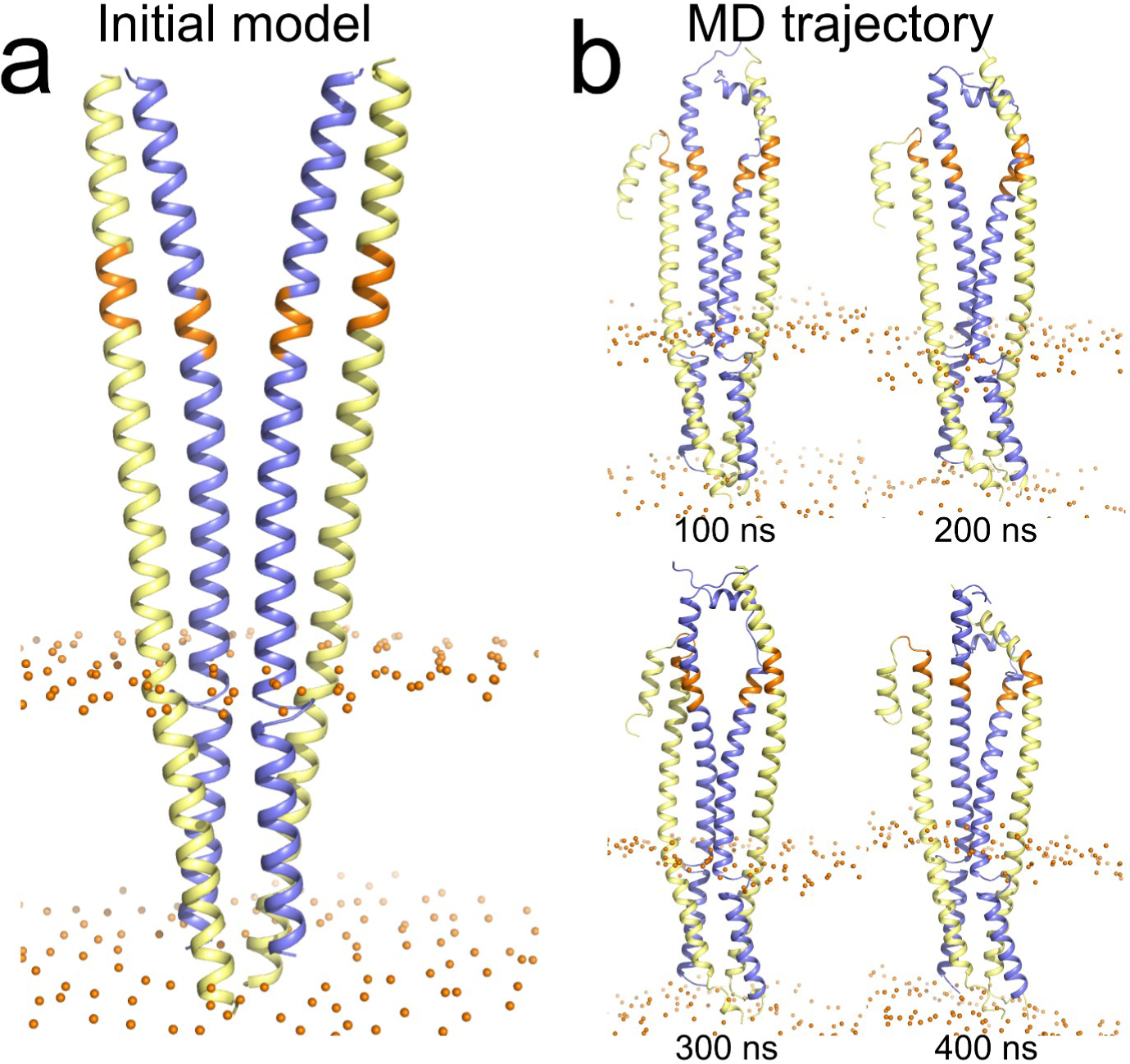
The Y-model of the FtsLB complex. a) Initial model. The model has the same **transmembrane** region of the original I-model. The coiled-coil region was modeled as a pair of two-stranded coiled-coil domains, each containing one FtsL and one FtsB helix. The modeling is based on the same set of side chain contacts between FtsB and FtsL inferred by the co-evolutionary analysis used to derive the I-model. Colored in orange is the CCD region. Spheres: lipid headgroup phosphate P atoms. b) Four frames of the trajectory of a 400 ns molecular dynamic run of the Y-model. The transmembrane and coiled-coil region remain relatively stable during the entire trajectory. The helices tend to break and occasionally unfold in correspondence of a predicted hinge near the CCD region (orange). The RMSD analysis of all three replica run is reported in Fig. S6 and in supplementary Table S3.

### AlphaFold2 modeling of FtsLB supports the structural features of the Y-model

With the outstanding performance of the program AlphaFold2^46^ at the recent CASP14 structural prediction competition, we decided to compare prediction of the FtsLB complex obtained with this method to our models. AlphaFold2 produced five models for FtsLB, which are ranked by their confidence score (supplementary Fig. S5 and Table S2). Although the program was given a 2:2 FtsL:FtsB stoichiometry as the input, only the fifth ranked model produced a tetramer. Interestingly, the fifth model assumed a configuration that roughly resembles the Y-model, with its tetrameric assembly being mediated by the transmembrane region and two separated two-stranded coiled-coil domains (Fig. S5a). However, rank model 5 and the Y-model align poorly (Cα RMSD of 11.11 Å). It should be noted that rank model 5 is a low-confidence model that is very loosely packed in its transmembrane region, and thus it is unlikely to be a good candidate for the structure of FtsLB.

The top four ranked AlphaFold2 models form two separated FtsB-FtsL heterodimers (Fig. S5a). This is contrary to our experimental evidence, which indicates that FtsLB forms a heterotetramer^28, 29^. In spite of this difference, however, ranked models 1-4 are structurally very similar to each half of the tetrameric Y-model (Cα RMSD values of 2.26-2.35 Å, Table S2), whereas their alignment with the I-model is less optimal (Cα RMSD values of 3.26-3.47 Å). The AlphaFold2 models also display the same distinctive structural features that we identified, with identical interfaces, an unwound loop in the Gly-rich linker, and FtsL in a continuous helical configuration through the transmembrane and coiled-coil domains. The superimposition of AlphaFold2 ranked model 1 against the Y-model in supplementary Fig. S5c clearly illustrates how the two models that are in excellent agreement. In summary, aside from the stoichiometry of the complex, the AlphaFold2 results provide a useful, independent validation of the structural organization of the Y-model.

### The Y-model remains more stable in comparison to the I-model during MD simulations

To assess the stability of the Y-model and its dynamic properties, we performed molecular dynamics (MD) simulations in explicit POPE bilayers in conditions analogous to the previous MD simulations of the I-model^29^. As done previously, the coiled coil was extended in the initial configuration by approximately 20 amino acids beyond its likely boundaries (the CCD region) to avoid end-effects. We will refer to this C-terminal region (residues 92-110 for FtsL and 62-79 for FtsB) as the post-CCD region. Three replica MD simulations were run for 400 ns each. The three trajectories are illustrated in Fig. 6b and, in more detail, in supplementary Fig. S6. Overall, we observed that the Y-model remained stable during the simulation time.

Similar to our previous simulations^29^, the transmembrane region underwent only minor rearrangements, with average RMSDs of 2.4, 1.6, and 1.8 Å in the three replica runs (Fig. S6, red traces, and supplementary Table S3). We found that the reconfigured coiled-coil region of the Y-model was very stable across all three replica runs with 1.4-1.7 Å average RMSDs (green and blue traces). For comparison, the average RMSDs for the same region of the I-model were 2.3-3.4 Å^29^. The fact that the coiled coil of the Y-model remained well structured is in stark contrast with the simulation of the I-model, during which the four-stranded coiled coil opened, allowing water molecules into the core and in contact with the polar cluster.

As hypothesized, the terminal polar moieties of the long side chains (the amide group of FtsB Gln-39 and the guanidinium group of FtsL Arg-67 and Arg-74) are partially solvent exposed in the two-stranded coiled coils of the Y-model, while their non-polar CH 2 groups contribute to hydrophobic packing at the interface. Solvent Accessible Surface Area (SASA) calculations indicate that on average the amide group of Gln-39 remains 20-40% solvent accessible and the guanidinium groups of Arg-67 and 74 remain 40-60% accessible. The only polar group that remained nearly completely buried in the coiled coil was the shorter Asn-43 of FtsB, but its hydrogen bonding potential is satisfied by interacting with the backbone carbonyl group of Leu-70 and often with the side chain of Glu-73 on the opposed FtsL chain. In fact, the four side chains of the polar cluster tend to interact with each other in a shared network of hydrogen bonds (Fig. 7b). Overall, the organization of the polar cluster appears clearly more favorable in the Y-model in comparison with the I-model, in which these positions were unfavorably buried in a four-helix coiled-coil conformation.

**Fig. 7.**
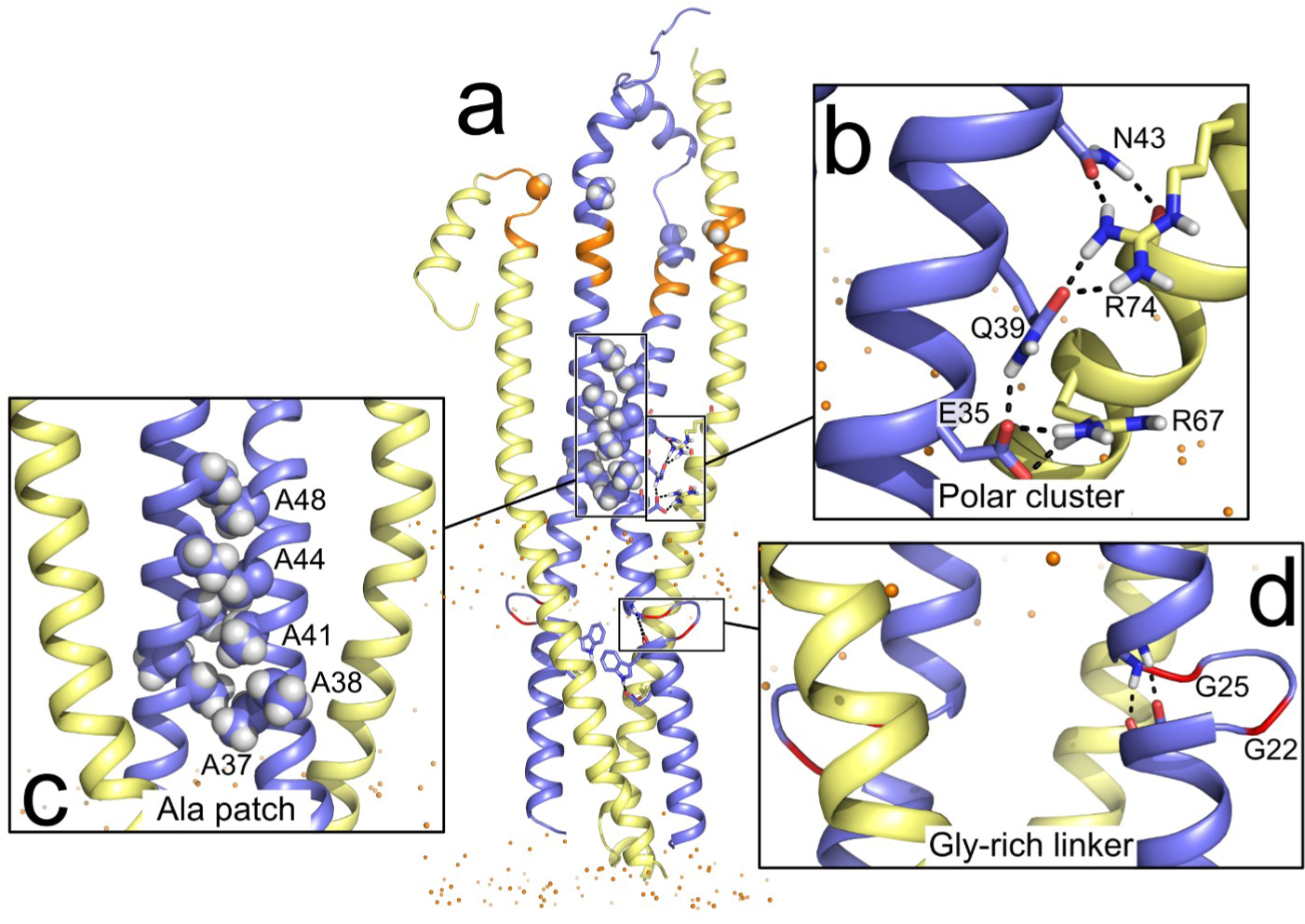
Features of the Y-model. a) MD frame of the Y-model highlighting three features of interest. b) The amide group of Gln-39 and the guanidinium group of Arg-67 and Arg-74 are partially solvent exposed whereas only the shorter Asn-43 is mainly buried. The polar cluster side chains tend to interact with each other in a shared network of hydrogen bonds. c) A patch of Ala residues on the opposite face of the FtsL/FtsB coiled-coil interfaces mediates the interaction of the two FtsB helices. The interaction is stable through the entire replica MD runs. d) The FtsB Gly-rich linker forms a short four-amino acid loop. The helical termini of the transmembrane and coiled-coil helices remain in contact in a configuration that resembles a continuous helix. There is hydrogen bond formation between the carbonyl groups of transmembrane residues 19-20 and the N–H groups of residues 26-27 of the coiled coil.

The coiled coils displayed a tendency to break near the CCD region, in correspondence to a likely hinge that contains Gly residues in both FtsB (positions 62 and 63) and FtsL (position 92). The terminal segments beyond this section are highly dynamic in our simulations, as indicated by their high RMSD traces (Fig. S6, orange and purple traces). This is primarily due to the lever arm effect, since most of the RMSD arises from propagation of the unfolding of the hinge region, while the helical portion past the hinge stays mostly helical. The post-CCD segment of FtsB is essential for binding to FtsQ^33, 47^ and is known to be structured in the FtsQLB complex^31, 32^. This segment forms an eleven-residue helix that starts right after the predicted hinge (position 64) and then associates with the C-terminal β-sheet of FtsQ by β-strand addition. FtsQ was not included in our simulations, and thus it is not surprising that the C-terminal peptides unfolded in the simulation, although significant helical content was generally retained. The structure of the C-terminal segment of FtsL is not known, although presumably this terminal tail is also structured in the presence of FtsQ (or other components). In our simulations, this segment also tends to extend at the predicted hinge but otherwise retains significant helical content, similar to the corresponding segment of FtsB.

### An Ala-rich patch in FtsB mediates inter-coil contact in the Y-model

An interesting outcome of the MD simulations is a small but significant rearrangement of the two branches of the “Y”. The two coiled-coil domains are separated by approximately 7 Å in the initial model, but this gap closes in the simulations. After this closure, the domains remain in permanent contact during all runs. Their contact is mediated by a mildly hydrophobic patch in FtsB consisting of five Ala residues that are clustered on the solvent-exposed, back side of the helix (positions 37, 38, 41, 44, and 48, Fig. 7c). This interaction results in an additional helix-helix interface with an ∼1,000 Å^2^ contact area spanning approximately three helical turns and a right-handed crossing angle of approximately -35°.

The interaction forms rapidly within a few nanoseconds in runs 1 and 2. Once the contact is established, it persists for the entirety of the 400 ns runs, and thus the two branches of the “Y” become locked into proximity. Only in replica run 3, the branches initially splay apart forming a more open “Y”, and it takes almost 40 ns for the branches to come in contact. In this run, when the contact is formed it is not symmetrical, involving Ala residue 41, 44, and 48 on one chain packing with residues 37, 41 and 44 in the other chain. Once this interaction is established, it also persists for the entirety of the 400 ns simulation, but it does not revert to the nearly symmetrical conformation observed in the other runs. Because of the asymmetry, one coil of FtsB appears to be pulled away from the transmembrane region, thus stretching the Gly-rich loop and breaking the contact between the termini of the transmembrane and coiled-coil helices.

To test whether the Ala patch is an essential feature of FtsLB, we drastically mutated the Ala residues and measured the resulting cell length phenotypes *in vivo*. All five Ala residues were mutated at once to combinations of Asp and Glu (AAAAA→EEEDD and AAAAA→DDDEE), according to the hypothesis that the replacement of hydrophobic Ala with negatively charged amino acids should destabilize the interaction interface. As shown in Fig. 8, the mutations did not cause elongation phenotypes, but their distributions were enriched in smaller cells compared to wild type (2.74 and 2.69 μm median length for the EEEDD and DDDEE mutants, respectively, compared to 2.93 μm for the wild type), in a fashion that is similar to the FtsL R74E mutant discussed previously.

**Fig. 8.**
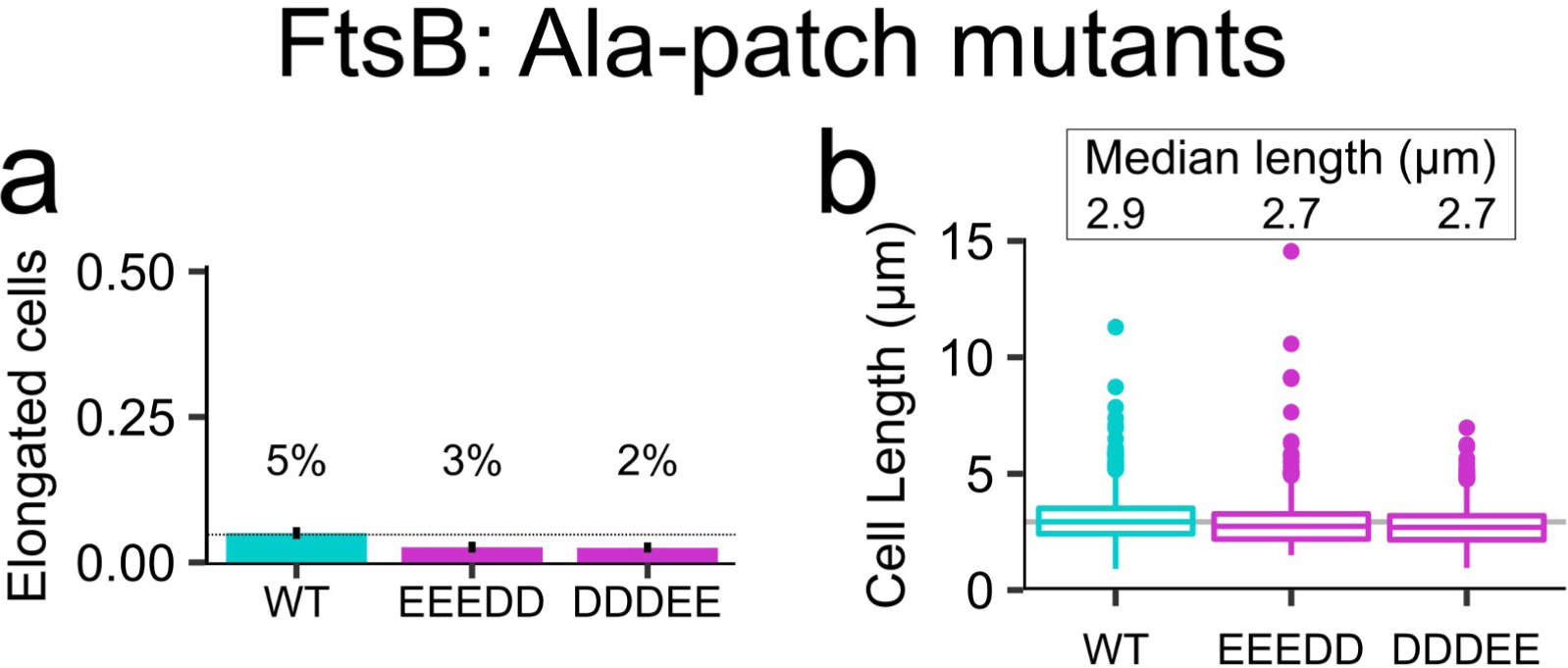
Disruption of the Ala-patch of FtsB results in smaller cells. A mildly hydrophobic patch of Ala residues is predicted to mediate the interaction of the two FtsB helices in the Y-model. All five Ala positions (37, 38, 41, 44, and 48) were simultaneously replaced with negatively charged residues, with a combination of three Glu and two Asp residues (EEEDD) and vice versa (DDDEE). a) Fraction of elongated cells for the mutants. Error bars represent the 95% confidence interval, estimated from 1000 replicates of bootstrap resampling. b) Median cell length distributions. The disruption of the potential interaction interface does not yield a defective division phenotype; however, a shift of the distribution with an increase of small cells suggests that the disruption of the Ala-patch induces some level of disregulated triggering of early cell division. Experiments were performed at 37 °C. Cell distributions are plotted in supplementary Fig. S2.

Perhaps surprisingly, our experiments indicate that the disruption of the Ala patch feature does not cause major loss of function. If the Y-model is an accurate representation of the FtsLB complex, the data indicate that the added stability deriving from the interaction of the two coiled-coil branches is not a strong requirement. However, the observed enrichment in small cells is consistent with the hypothesis that fine-tuning the stability of the coiled coil participates in governing the balance between the *on* and *off* states of the complex. Specifically, it suggests that destabilization of the coiled coil tends to favor unregulated (early) cell division. It remains to be confirmed whether the hypothesized destabilization arises from the loss of the inter-branch hydrophobic interface or from direct destabilization of the coiled coil due to the presence of a patch of negative residues.

### The AWI positions of FtsL are surface-accessible in an initial model of the FtsQLB complex

The CCD and AWI positions are located in a region near the end of the predicted coiled coil in FtsLB, preceding a likely hinge created by three Gly residues in close proximity (positions 62-63 in FtsB and 92 in FtsL). CCD mutants rescue a *ΔftsN* phenotype, suggesting that they induce changes in the FtsLB complex that mimic the activation signal given by FtsN^21, 24^. This process likely involves a conformational change that makes the AWI positions of FtsL available for interacting with FtsI, leading to activation of septal PG synthesis^25, 26^.

In the Y-model, the AWI positions (82-84, 86-87, and 90) of FtsL are completely solvent exposed. Given that FtsLB exists as a complex with FtsQ, we assembled a preliminary model of the FtsQLB complex by aligning the region of FtsB that is in common between the Y-model and the co-crystal structure of the C-terminal fragment in complex with FtsQ (PDB code 6H9O^32^). This fragment starts with the post-CCD helix (residues 64-74), following the putative hinge at the end of the coiled coil (62-63). Because of the flexible hinge, the orientation of the post-CCD helix is uncertain. However, the N-terminus of the periplasmic domain of FtsQ needs to be near the lipid bilayer for proper placement of its transmembrane helix. We found that by modeling the post-CCD helix as a nearly straight continuation of the coiled-coil helix, this constraint is satisfied and FtsQ is oriented in a position that does not collide with the FtsLB complex.

In this preliminary structural model of the FtsQLB complex (illustrated in Fig. 9a and b, with FtsQ depicted in gray), the surface of the coiled-coil region of FtsB is almost completely occluded. The helices of FtsB are flanked on one side by FtsL and on the opposite side by the helix of FtsB from the opposite coiled coil, and their remaining exposed face is then occluded by FtsQ. Conversely, the helix of FtsL packs only with its partner FtsB helix while the opposite helical face remains exposed to solvent even in the presence of FtsQ. Interestingly, all amino acids of the AWI region of FtsL (in green) occur in this face, and thus remain solvent exposed. This configuration supports the AWI region as a good candidate surface for interaction with FtsWI. Conversely, the coiled coil of FtsB, which is nearly completely buried in the Y-model, would require major structural rearrangements to become available.

**Fig. 9.**
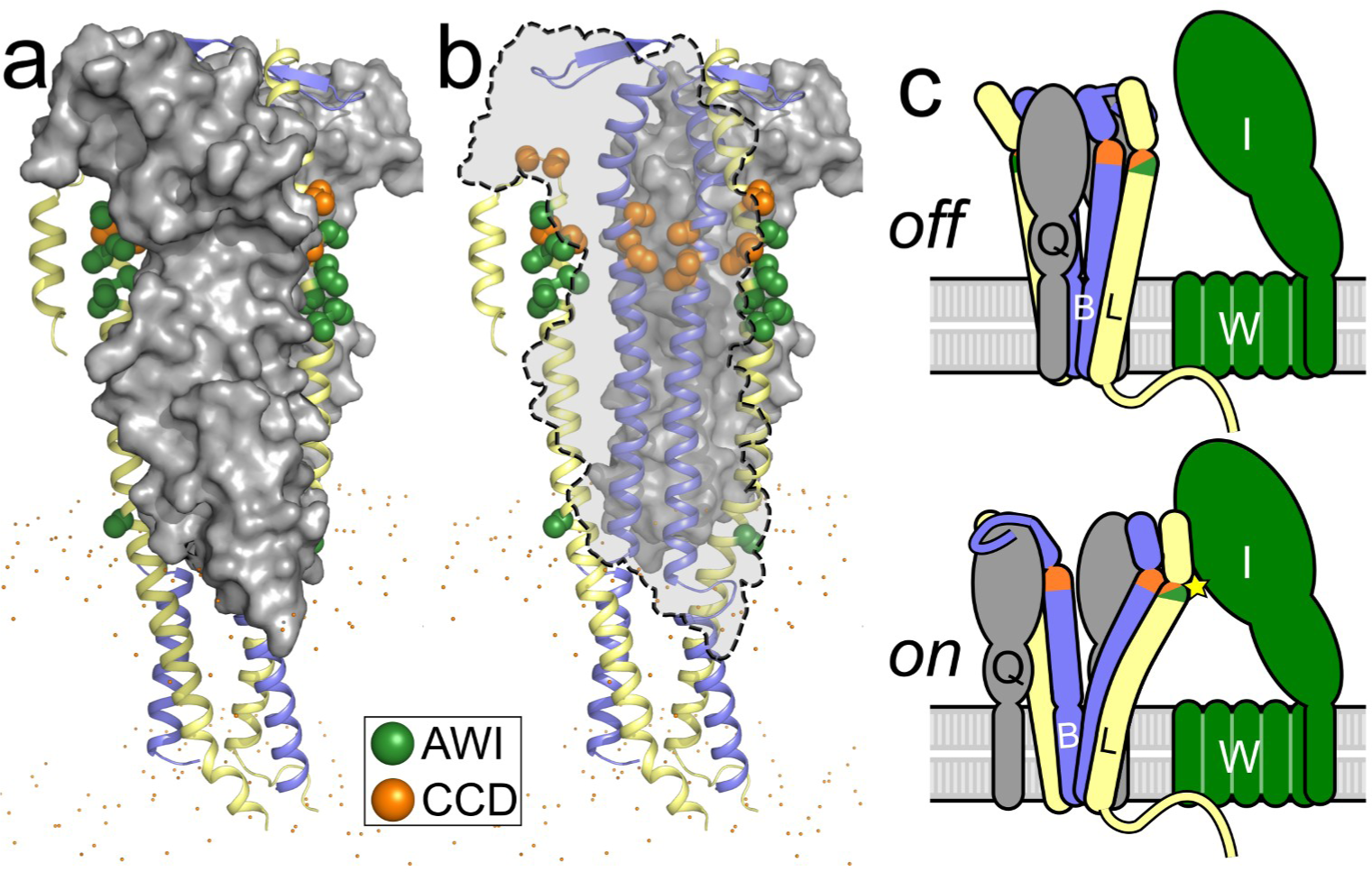
A structural shift in the coiled coil of FtsLB may be involved in its activation. *a*) Preliminary model of the FtsQLB complex, obtained by aligning the region of FtsB that is in common between the Y-model and the crystal structure of FtsQ (pdb 6H9O). In this model, the four helices of the coiled coil of FtsLB form a flat bundle sandwiched between the two periplasmic domains of FtsQ (grey). *b*) See-through representation of the same model, with one of the two FtsQ subunits removed and represented by a dashed outline. The positions of the CCD and AWI domains of FtsB and FtsL are highlighted as orange and green spheres, respectively. FtsB is nearly completely buried. FtsL is more solvent exposed and its AWI positions occur on a solvent-facing surface. *c*) Schematic representation of a potential model for FtsLB activation. A structural shift involving the coiled coil of FtsLB (potentially involving the separation of the two branches) exposes the AWI region of FtsL to a hypothesized protein-protein interaction with FtsI (star). This transition may be favored by destabilization of the coiled-coil region.

## Conclusions

In this article, we address the hypothesis that the coiled coil of FtsLB is a critical functional element. We explore the role of an unusual cluster of polar amino acids that occur at core positions within this domain and investigate whether the stability of the coiled coil is marginal by design. We propose that this characteristic may be critical for enabling the hypothesized FtsLB *off/on* transition that leads to the activation of the PG synthase machinery. Our evidence indicates that mutations affecting the conserved polar cluster lead to appreciable division phenotypes, indicating that this feature is functionally important. The fact that mutations to hydrophobic residues increase thermal stability *in vitro* but often lead to division defects *in vivo* suggests that the coiled-coil domain is indeed likely to be a “detuned” functional element, sacrificing some of the stability of the complex in order to enable proper function.

In addition to affecting the plasticity of the coiled coil, the polar cluster likely contributes to the shape of the periplasmic domain by causing it to branch out, as proposed in the revised Y-model of the complex. The branched Y-model satisfies all the known constraints (i.e., oligomeric state, inferred amino acid contacts, and mutagenesis data) as well as the original I-model, but it appears to be structurally more stable when assessed by molecular dynamics, unlike the I-model in which the coiled coil was rapidly infiltrated by water during its simulation^29^. The structural arrangement of the Y-model is also in better agreement with predictions from AlphaFold2, after accounting for discrepancies between their overall stoichiometries. For these reasons, we propose the Y-model as the more likely configuration of the FtsLB complex.

This hypothesis that the polar cluster shapes the coiled coil into independent branches offers an interpretation for our observation *in vitro* that the “idealized” coiled coil (the “4×-mutant” FtsBQ_39L,N43I_/FtsL_R67I,R74I_) unfolds at higher temperatures but displays a broad non-cooperative transition – an indication that the complex may assume multiple alternative configurations. In the “idealized” mutant, the coiled coil would not pay a steep penalty (i.e., the burial of polar side chains) to continue in the same four-stranded configuration of the transmembrane region, thus it is possible that a competition could occur between the I- and Y-configurations.

One of the most interesting outcomes of this analysis is the discovery of suppressor mutations that occur within the polar cluster. Specifically, changes in FtsL at position Arg-74 can suppress the phenotype of mutations at position Arg-67. The observations suggest a likely structural interplay between the two positions, where mutations cancel each other’s opposing tendencies to make division less likely (R67I and R67E, which display moderate elongation) or more likely (R74E, which is enriched in smaller cells), suggesting that these positions fine-tune the interactions that govern the balance between the *on* and *off* states of the complex. In addition, we noted that disruption of the mildly hydrophobic Ala-patch (EEEDD and DDDEE mutants) that mediates the contact between the two branches of the coiled coil in the Y-model MD simulations also results in a smaller than normal cell distribution. This finding suggests that separation of the branches, which would be favored in the EEEDD and DDDEE mutants, facilitates the *on* transition. It is unclear if the R74E mutation operates in a similar manner, although in the model, substitution of R74 disrupts an intricate network of hydrogen bonding that connects the entire polar cluster, as illustrated in Fig. 7b.

We propose a model in which the stability of the FtsLB coiled coil needs to be perfectly tuned. If the coiled coil is too stable, the complex does not respond appropriately to the FtsN- derived stimulus that normally causes activation of the complex. Conversely, if the coiled coil is further destabilized, FtsLB can become activated in an unregulated manner. Functional, metastable coiled-coil domains are known to be critical for other bacterial signaling processes such as those involving two-component systems^48^, in which the coiled coil functionally connects the membrane-bound sensory domain with the response kinase domain in the cytoplasm. Specific examples include the *Staphylococcus aureus* antibiotic sensor NsaS^49^, the *Bacillus subtilis* thermosensor DesK^50, 51^, and the *Bordetella pertussis* virulence factor BvgS^52^, each of which contains a coiled-coil domain with nonideal residues that enable conformational changes needed to regulate signaling activity. For these systems, the degree of hydrophobicity of the coiled-coil interface impacts the rigidity of the sensor and can even directly modulate the balance between kinase and phosphatase activity^51, 52^. The presence of nonideal residues at the interface of the FtsLB coiled coil suggests that a similar “tuning” of stability may be needed to support conformational rearrangements.

Further structural and biophysical characterization is necessary to validate the model and to fully understand how the structural organization of the coiled coil and its conformational changes operate in triggering septal PG reconstruction.

### Experimental Procedures

#### Plasmid cloning

For the *in vivo* complementation experiments, mutant variants of FtsB or FtsL were cloned via standard QuikChange mutagenesis or inverse PCR into pMDG7^47^ (flag3-FtsB) or pMDG29^30^ (flag3-FtsL), respectively. For the CD experiments, the His-tagged FtsB and Strep-tagged Cys-less (C41A and C45A) FtsL35–121 were ligated into a modified pETDuet-1 vector at restriction sites NcoI/HindIII and NdeI/XhoI, respectively. Point mutations were introduced using standard QuikChange mutagenesis. All constructs were confirmed by DNA sequencing (Quintara Biosciences). A complete plasmid inventory is included in supplementary Table S4.

#### Bacterial strains, plasmids, and media for in vivo experiments

The phenotypic analyses were performed using depletion strains NB946^53^ for FtsB and MDG277^47^ for FtsL (both obtained from Jon Beckwith and associates) in which the WT copy of the protein of interest is under control of a repressible PBAD promoter within the chromosome. These strains were transformed with plasmids containing either WT protein (positive control), empty vector (negative control), or a mutant version of the protein to test for defects in cell division as evidenced by an increase in cell length. For all experiments described, bacterial cells were grown in LB medium supplemented with 100 μg/mL spectinomycin (Dot Scientific) and the appropriate carbon source. Medium was supplemented with 0.2% (w/v) L-arabinose (Sigma) or 0.2% (w/v) D-glucose (Sigma) to induce or repress, respectively, the expression of chromosomal copies of the WT genes regulated by the PBAD promoter. 20 μM isopropyl-β-D-1-thiogalactoside (IPTG) was added to the medium to induce the expression of mutant genes regulated by the Ptrc promoter in the plasmid.

#### Depletion strain experiments

The protocol for the depletion strain experiments was adapted from Gonzalez and Beckwith^47^. In short, a mutated copy of FtsB or FtsL was transformed into its respective depletion strain. Strains were grown overnight at 37 °C on an LB plate supplemented with arabinose and spectinomycin. A single colony from the plate was grown overnight at 37 °C in 3 mL of LB medium supplemented with arabinose and spectinomycin. The overnight culture was then diluted 1:100 into fresh LB medium containing the same supplements and grown to an OD_600_ of ∼0.3. An aliquot of 1 mL of culture was washed twice with LB medium lacking any sugar and then diluted 1:100 into 3 mL of fresh LB medium supplemented with glucose, IPTG, and spectinomycin to induce expression of the mutated gene in the plasmid and to repress the WT gene in the chromosome. The cells were then grown at 37 °C for 3.5 hr, the approximate time necessary to deplete the cells of the WT chromosomal copy^47^. The cells were then placed on ice to stop growth before imaging. Depletion strains provided with a WT copy of their respective protein in the plasmid were tested as positive controls, and, similarly, depletion strains with no protein in the plasmid (empty vector) were tested as negative controls.

#### Microscopy and cell length measurements

10 μL of cell samples were mounted on a number 1.5, 24 X 50 mm (0.16 – 0.19 mm thickness) cover glass slide (Fisher or VWR). Cells were cushioned with a 3% (w/v) agarose gel pad to restrict the movement of the live cells. Cells were optically imaged using a Nikon Eclipse Ti inverted microscope equipped with crossed polarizers and a Photometrics CoolSNAP HQ2 CCD camera using a Nikon X100 oil objective lens. Phase-contrast images of bacterial cells were recorded with a 50 ms exposure time using Nikon NIS Elements software. Multiple snapshots were collected for each experiment. All images were analyzed to measure the cell length in Oufti^54^ using one single optimized parameter set and manual verification. Confidence intervals of the fraction of elongated cells for each mutant were computed from 1000 replicates of bootstrap resampling^55^.

#### Western blots

Expression level across all variants was assessed by Western blot analysis (supplementary Fig. S3). 3.0 mL of cells were pelleted and resuspended in 300 μL of lysis buffer (50 mM HEPES pH 8.0, 50 mM NaCl) with 5 mM β-Mercaptoethanol (βME). The cells were sonicated and centrifuged at 21,000 × g for 10 min before collecting the supernatant. Total protein concentration was determined by BCA assay (Pierce). 120 μL of lysates were mixed with 40 μL of 4× LDS sample buffer (Novex, Life Technologies) with βME and boiled at 98 °C for 3 min. For each FtsL or FtsB sample, the equivalent of 7 μg or 15 μg, respectively, of total protein was separated by SDS-PAGE (Invitrogen) and transferred to polyvinylidene difluoride membrane (VWR). Horseradish peroxidase-tagged anti-FLAG (M2) antibodies (Sigma; 1:1,000) were used for immunoblotting analysis.

#### Protein expression and purification for CD

Plasmids were transformed into BL21(DE3) cells (NEB) and plated overnight at 37 °C on on LB agar with 100 μg/mL ampicillin. Cells were washed off the plates with 1 mL LB broth and inoculated into 1 L of ZYP-5052 autoinduction medium as described^56^ and grown at 37°C until reaching an OD_600_ of ∼0.8, after which they were incubated overnight at 22 °C. Following expression, cells were pelleted, resuspended in cell wash buffer (100 mM NaCl, 10 mM HEPES pH 8.0), pelleted again, flash frozen, and stored at -80 °C for future use. The cells were then lysed by sonication in 10 mL/g lysis buffer (50 mM NaCl, 50 mM HEPES pH 8.0) supplemented with 0.5 mg/mL lysozyme, 5 mM β-Mercaptoethanol (βME), 1 mM phenylmethylsulfonyl fluoride, 1 mM EDTA, and a protease inhibitor cocktail providing (final concentrations) 8 μM leupeptin (Peptides International), 11.2 μM E-64 (Peptides International), 0.32 μM aprotinin (ProSpec), and 0.32 mM 4-(2-aminoethyl)benzenesulfonyl fluoride (Gold BioTechnology). The inclusion body fraction was separated by centrifugation at 10,000 × g for 20 min, followed by ultracentrifugation of the supernatant at 180,000 × g for 30 min to isolate the cell membranes. The FtsLB complex was then extracted from the membrane fraction with lysis buffer supplemented with 18 mM n-decyl-β-D-maltopyranoside (DM; Anatrace) and 5 mM βME, rocking at room temperature overnight. Non-resuspended debris was separated from the solubilized protein via centrifugation at 10,000 × g for 20 min. The supernatant was added to ∼3 mL of Ni-NTA-agarose resin (Qiagen) and rocked for 2 hr at 4 °C for batch binding before performing gravity-flow purification. Purification was performed by running 10 column volumes of Ni wash buffer (300 mM NaCl, 25 mM HEPES pH 8.0, 50 mM imidazole, 1 mM βME) supplemented with 510 μM n-dodecyl-β-D-maltopyranoside (DDM; Avanti Polar Lipids) and 10 column volumes of elution buffer (300 mM NaCl, 25 mM HEPES pH 8.0, 300 mM imidazole, 1 mM βME) also supplemented with 510 μM DDM. Protein purity was assessed via SDS-PAGE (Invitrogen).

#### CD experiments

Purified FtsLB protein was dialyzed twice at room temperature for at least 2 hr into 1 L CD buffer (10 mM phosphate buffer pH 7.4, 100 mM NaF) supplemented with 170 μM DDM (1× critical micelle concentration to prevent detergent exchange), then overnight at 4 °C in 1 L CD buffer supplemented with 510 μM DDM. Samples were kept at 4 °C or on ice from this point forward. Protein concentration was determined against the final dialysis buffer using *A280* and an extinction coefficient of 32,430 M^-^cm^-^ for the FtsLB complex (calculated via ExPASy). Protein was diluted to ∼14 μM, then filtered with 0.22 μm (13 mm diameter) PVDF syringe filters (CELLTREAT) before redetermining the final protein concentration. Samples were degassed in a vacuum chamber for at least 30 min, then centrifuged for 20 min at 21,000 × g. The final dialysis buffer was also filtered and degassed in the same manner to use as a blank in the CD experiments. CD spectra were obtained using an Aviv model 420 CD spectrometer and quartz cuvettes with a 0.1 cm pathlength. All spectra were recorded in 1 nm increments, with either a 10 s or 20 s averaging time, and after a 5 min equilibration time upon reaching a 0.3 °C deadband. The spectra were baseline corrected by buffer subtraction. For the CD-monitored thermal melting experiments, the samples were heated at 3 °C intervals with a 20 s equilibration time. Because the transitions were not reversible, detailed thermodynamic analyses were not carried out, and the curves were only fitted to sigmoidal transitions to calculate their temperature midpoints (T_m_).

#### Bioinformatic analysis

Homologues of FtsB and FtsL were collected using the DELTA-BLAST algorithm^57^ on the RefSeq database^58^. FtsB-FtsL pairs were selected by the NCBI taxonomic identifier. In the case of multiple sequences per taxa, the one with the lowest E-value to the query *E. coli* FtsB or FtsL sequence was selected. Proteobacterial sequences were identified via the NCBI taxonomy database^59^. Sequences were aligned using the MAFFT algorithm^60^. Statistical analyses were performed in R^61^ with the aid of the following packages: tidyverse^62^, tidymodels^63^, Biostrings^64^, zoo^65^, taxize^66^, rentrez^67^, and tidygraph^68^.

### Molecular modeling

Modeling of the rearranged FtsLB complex was performed as described previously ^29^. Briefly, the FtsLB heterodimer was modeled using a Monte Carlo procedure to model supercoiled helical bundles^69^. The superhelical radius (*r*_1_), superhelical pitch (*P*), helical rotation (Φ_1_), and z-shift (*s*) of both FtsL_52-94_ and FtsB_21-63_ were freely altered, whereas the rise per residue (*h*) and helical radius (*r*_0_) were kept constant. Energies were calculated based on CHARMM 22 van der Waals and CHARMM 22 electrostatic terms with additional sigmoidal distance restraints for each pair of evolutionary couplings in the coiled-coil region^29^. The heterodimeric FtsLB coiled coil was then aligned with one half of the previously modeled heterotetrameric transmembrane domain using residues 52-58 of FtsL, which were present in both models. Both domains were kept parallel to the Z-axis. The juxtamembrane regions of FtsL and FtsB were then replaced with loops corresponding to fragments from the PDB. For FtsB, six-residue loops (corresponding to positions 21–26) with four flanking helical residues on each side were used, with an additional sequence requirement that the fragment contain at least one glycine. For FtsL, 15-residue fragments with four flanking helical residues on each side were used with the requirement that the loop have helical secondary structure. This arrangement was made C2-symmetric to generate the Y-model. Finally, the side chains were repacked using a greedy trials algorithm, and the model was minimized using BFGS constrained optimization in CHARMM^70^.

### Protein structure prediction using AlphaFold2

The FtsLB complex was predicted with AlphaFold using the AlphaFold2_advanced notebook from Colabfold^46, 71^, which allows for predictions of protein complexes. The full sequences of FtsB and FtsL were used as input, with two chains for each protein to model a 2:2 heterotetramer. Mmseqs2^72^ was chosen for MSA generation, and the use_turbo option was enabled.

### All-atom molecular dynamic simulations

For the molecular dynamics simulations, the model’s coiled-coil region was extended to avoid end-effects to residues 110 (FtsL) and 79 (FtsB). The cytoplasmic side of FtsL was also extended to include residues 30-34, modeled in ideal α-helix. Three 400 ns all-atom MD simulations were performed using the CHARMM36m force field^73^ and NAMD 2.10 software^74, 75^. CHARMM-GUI membrane builder^76^ was used to prepare systems composed of a POPE bilayer consisting of 301 lipids, the FtsLB tetramer, an ionic concentration of 0.150 M NaCl, and 59163 TIP3P water molecules for hydration. The size of the boxes at the beginning of the simulation were approximately 97 × 97 × 242 Å^3^. The simulations were initially minimized and equilibrated for 75 ps at an integration time of 1 fs/step and for 600 ps at an integration time of 2 fs/step. The integration time step for the production runs of each of the systems was 2.0 fs/step. The simulations were carried out in the NPT ensemble at a pressure of 1 atmosphere and a temperature of 310.15 K, using the Nose-Hoover Langevin piston and Langevin dynamics method. Particle Mesh Ewald was used for electrostatic interactions, and a 12 Å cutoff was applied to Lennard-Jones interactions with a switching function from 10 to 12 Å. The RMSD analysis was performed using the RMSD trajectory tool in VMD^77^. Hydrogen bonding analysis was performed with an in-house script.

## Acknowledgments

This work was supported in part by National Institutes of Health (NIH) Grant R01-GM099752 and R35-GM130339 to A.S. and NSF Grant CHE-1829555 to Q.C. S.J.C. was supported in part by a Dr. James Chieh-Hsia Mao Wisconsin Distinguished Fellowship. S.G.F.C. was supported in part by the Arthur B. Michael Departmental Fellowship and the William R. & Dorothy E. Sullivan Wisconsin Distinguished Graduate Fellowship. G.D.V. was supported in part by a SciMed GRS Fellowship.

## Conflicts of Interest

The authors declare that they have no conflicts of interest with the contents of this article.

## Data availability

Bacterial strains and plasmids are available from the authors upon request. The Y-model and the AlphaFold2 model of FtsLB can be downloaded from http://seneslab.org or from the authors upon request. The MSL libraries are available for download from https://sourceforge.net/projects/mslib/. Code is also available from the authors upon request.

## SUPPLEMENTARY INFORMATION

### Supplementary tables

**Table S1.** Frequency of polar amino acids at a and d positions in a database of 2,662 crystal structures of coiled coils from the CC+ database.

| **Two-stranded parallel coiled coils** |  |  |  |
| --- | --- | --- | --- |
| Amino acid | *a* positions<sup>1</sup> | *d* positions<sup>1</sup> | *a+d* positions<sup>1</sup> |
| D | 0.97% | 1.13% | 1.05% |
| E | 3.34% | 4.26% | 3.80% |
| H | 1.79% | 1.46% | 1.63% |
| K | 4.72% | 2.79% | 3.76% |
| N | 3.22% | 1.72% | 2.47% |
| Q | 3.45% | 2.57% | 3.01% |
| R | 4.47% | 1.59% | 3.03% |
| **Total polar** | **21.97%** | **15.53%** | **18.76%** |
| **Three-stranded parallel coiled coils** |  |  |  |
| Amino acid | *a* positions<sup>1</sup> | *d* positions<sup>1</sup> | *a+d* positions<sup>1</sup> |
| D | 0.38% | 0.76% | 0.57% |
| E | 2.05% | 1.39% | 1.72% |
| H | 1.15% | 1.46% | 1.31% |
| K | 0.83% | 0.95% | 0.89% |
| N | 1.86% | 4.50% | 3.19% |
| Q | 5.00% | 5.26% | 5.13% |
| R | 1.09% | 1.46% | 1.27% |
| **Total polar** | **12.38%** | **15.77%** | **14.09%** |
| **Four-stranded parallel coiled coils** |  |  |  |
| Amino acid | *a* positions<sup>1</sup> | *d* positions<sup>1</sup> | *a+d* positions<sup>1</sup> |
| D | 0.16% | 1.20% | 0.69% |
| E | 2.86% | 2.07% | 2.46% |
| H | 1.14% | 1.59% | 1.37% |
| K | 1.14% | 0.48% | 0.81% |
| N | 1.39% | 1.75% | 1.57% |
| Q | 2.45% | 5.18% | 3.83% |
| R | 0.49% | 1.43% | 0.97% |
| **Total polar** | **9.63%** | **13.71%** | **11.69%** |
| **All parallel coiled coils** |  |  |  |
| Amino acid | *a* positions<sup>1</sup> | *d* positions<sup>1</sup> | *a+d* positions<sup>1</sup> |
| D | 0.84% | 1.08% | 0.96% |
| E | 3.14% | 3.77% | 3.46% |
| H | 1.69% | 1.45% | 1.57% |
| K | 4.00% | 2.38% | 3.19% |
| N | 2.93% | 1.99% | 2.46% |
| Q | 3.50% | 3.10% | 3.30% |
| R | 3.77% | 1.54% | 2.65% |
| **Total polar** | **19.87%** | **15.32%** | **17.60%** |

<sup>1</sup>Pseudocount-Adjusted frequency

**Table S2.** Alignment scores (Cα RMSD) of the five AlphaFold2 dimeric FtsLB models against half of the Y- and I-models (FtsBA 1-60, FtsLC 40-91)

| AlphaFold2<br>Model | pLDDT<sup>1</sup> | Y-model<br>Cα RMSD (Å) | I-model<br>Cα RMSD (Å) |
| --- | --- | --- | --- |
| Rank_1 | 79.78 | 2.33 | 3.47 |
| Rank_2 | 78.12 | 2.26 | 3.40 |
| Rank_3 | 77.63 | 2.27 | 3.36 |
| Rank_4 | 76.35 | 2.35 | 3.26 |
| Rank_5 | 61.94 | 2.71 | 2.31 |

<sup>1</sup>pLDDT score (predicted Local Distance Difference Test): AlphaFold2's overall confidence metric of the model (100 = most confident, 0 = least confident)

**Table S3.**
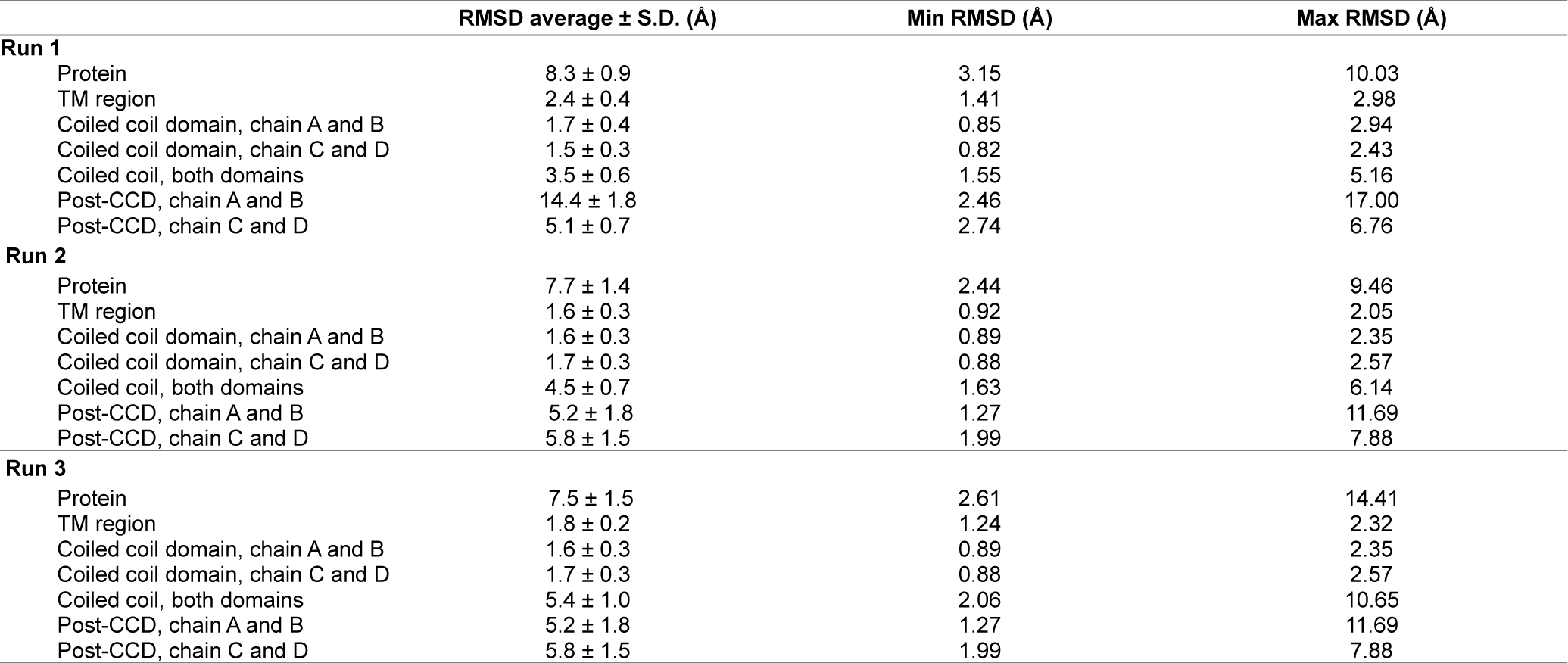
RMSD analysis of the three replica molecular dynamic runs of the FtsLB complex in the Y-model configuration.

**Table S4.** Strains and plasmids used in this work.

| **Strain/**<br>**plasmid** | **Description** | **Parent**<br>**vector** | **Source** |
| --- | --- | --- | --- |
| BL21(DE3) | chemically competent *E. coli* for protein overexpression | - | New England BioLabs (C2527) |
| NB946 | FtsB depletion strain | - | Buddelmeijer et al., 2002<sup>1</sup> |
| MDG277 | FtsL depletion strain | - | Gonzalez & Beckwith, 2009<sup>2</sup> |
| pSJC020 | His-FtsB \| Strep-FtsL35-121 C41A/C45A (Cys-less FtsL) | pETDuet-1 | Condon et al., 2018<sup>3</sup> |
| pSJC309 | His-FtsB Q39L/N43I \| Strep-FtsL35-121 C41A/C45A/R67I/R74I | pETDuet-1 | This paper |
| pNG162 | IPTG-inducible, low-copy-number vector (empty) | pAM238 | Goehring et al., 2006<sup>4</sup> |
| pMDG7 | flag3-FtsB | pNG162 | Gonzalez & Beckwith, 2009<sup>2</sup> |
| pSJC187 | flag3-FtsB Q39L | pMDG7 | This paper |
| pSJC188 | flag3-FtsB N43I | pMDG7 | This paper |
| pSJC208 | flag3-FtsL N50I | pMDG7 | This paper |
| pSJC287 | flag3-FtsB A37D/A38D/A41D/A44E/A48E | pMDG7 | This paper |
| pSJC288 | flag3-FtsB A37E/A38E/A41E/A44D/A48D | pMDG7 | This paper |
| pMDG29 | flag3-FtsL | pNG162 | Gonzalez et al., 2010<sup>5</sup> |
| pSJC183 | flag3-FtsL R67I | pMDG29 | This paper |
| pSJC185 | flag3-FtsL R74I | pMDG29 | This paper |
| pSJC190 | flag3-FtsL R67I/R74I | pMDG29 | This paper |
| pSJC194 | flag3-FtsL R67E | pMDG29 | This paper |
| pSJC201 | flag3-FtsL R74E | pMDG29 | This paper |
| pSJC220 | flag3-FtsL W81I | pMDG29 | This paper |
| pSJC254 | flag3-FtsL R67E/R74E | pMDG29 | This paper |
| pSJC304 | flag3-FtsL R67I/R74E | pMDG29 | This paper |
| pSJC324 | flag3-FtsL R67K | pMDG29 | This paper |
| pSJC325 | flag3-FtsL R74K | pMDG29 | This paper |
| pSJC326 | flag3-FtsL R67E/R74I | pMDG29 | This paper |
| pSJC334 | flag3-FtsL R74E/W81I | pMDG29 | This paper |
| pSJC335 | flag3-FtsL R74K/W81I | pMDG29 | This paper |
| pSJC336 | flag3-FtsL R67I/R74K | pMDG29 | This paper |
| pSJC337 | flag3-FtsL R67E/R74K | pMDG29 | This paper |

## Supplementary figures

**Fig. S1.**
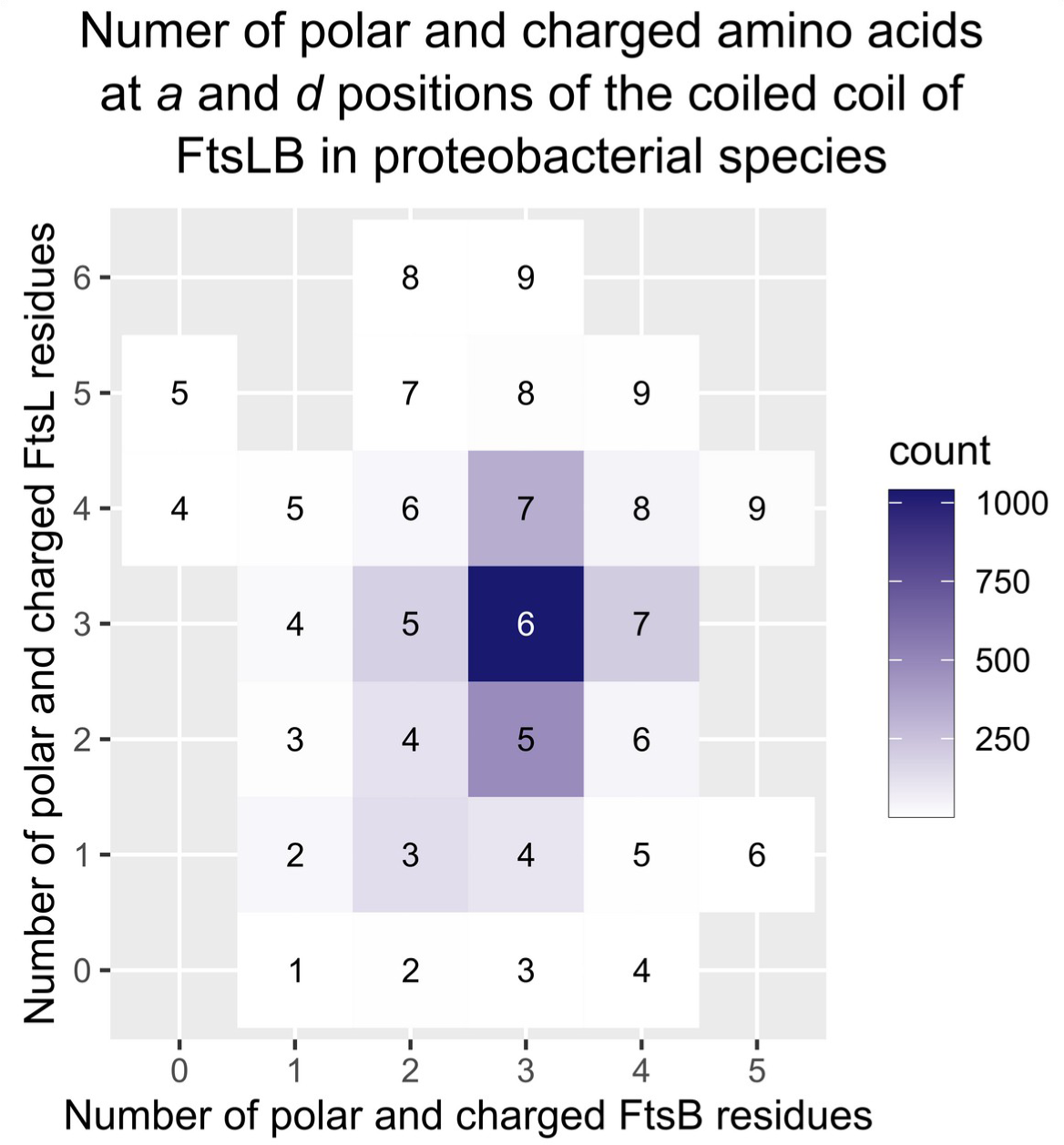
Conservation of the polar cluster in the core “*a*” and “*d*” positions of the coiled coil of FtsLB in proteobacterial species. Number of sequences with a given number of polar/charged amino acids (Asp, Glu, His, Arg, Lys, Gln, and Asn) contributed by FtsB (X axis) and FtsL (Y axis) at “*a*” and “*d*” positions in the five heptad repeats of the coiled coil (their sum is reported in the box). Data from an alignment of 2900 paired FtsB/FtsL proteobacterial sequences. The most frequent combination corresponds, by a large margin, to three polar residues contributed by both FtsB and FtsL, for a total of six polar residues. Although less frequent, combinations with a total of five or seven polar residues also occur relatively often. Overall, 84% of the sequences contain at least 5 polar residues.

**Fig. S2.**
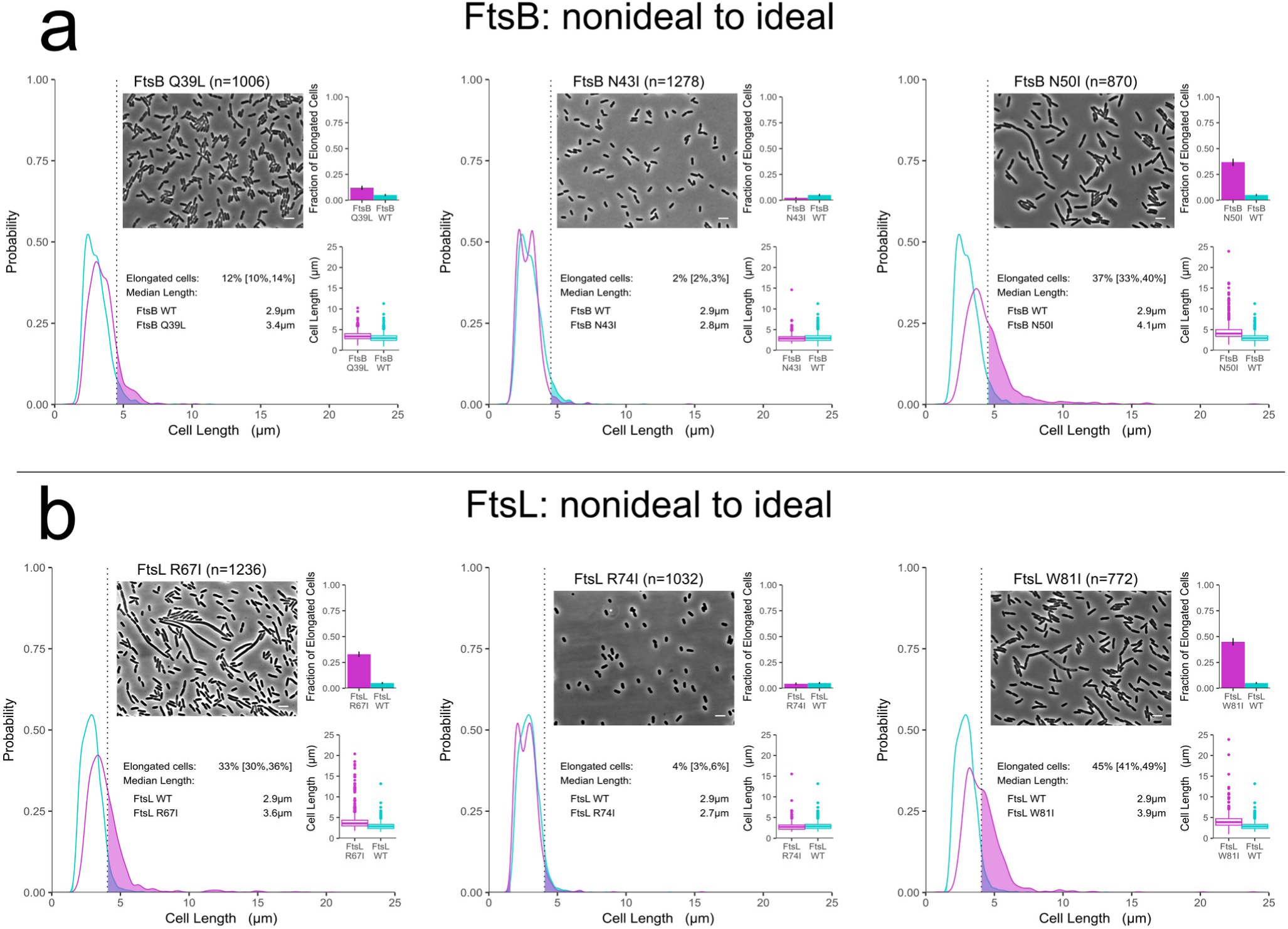
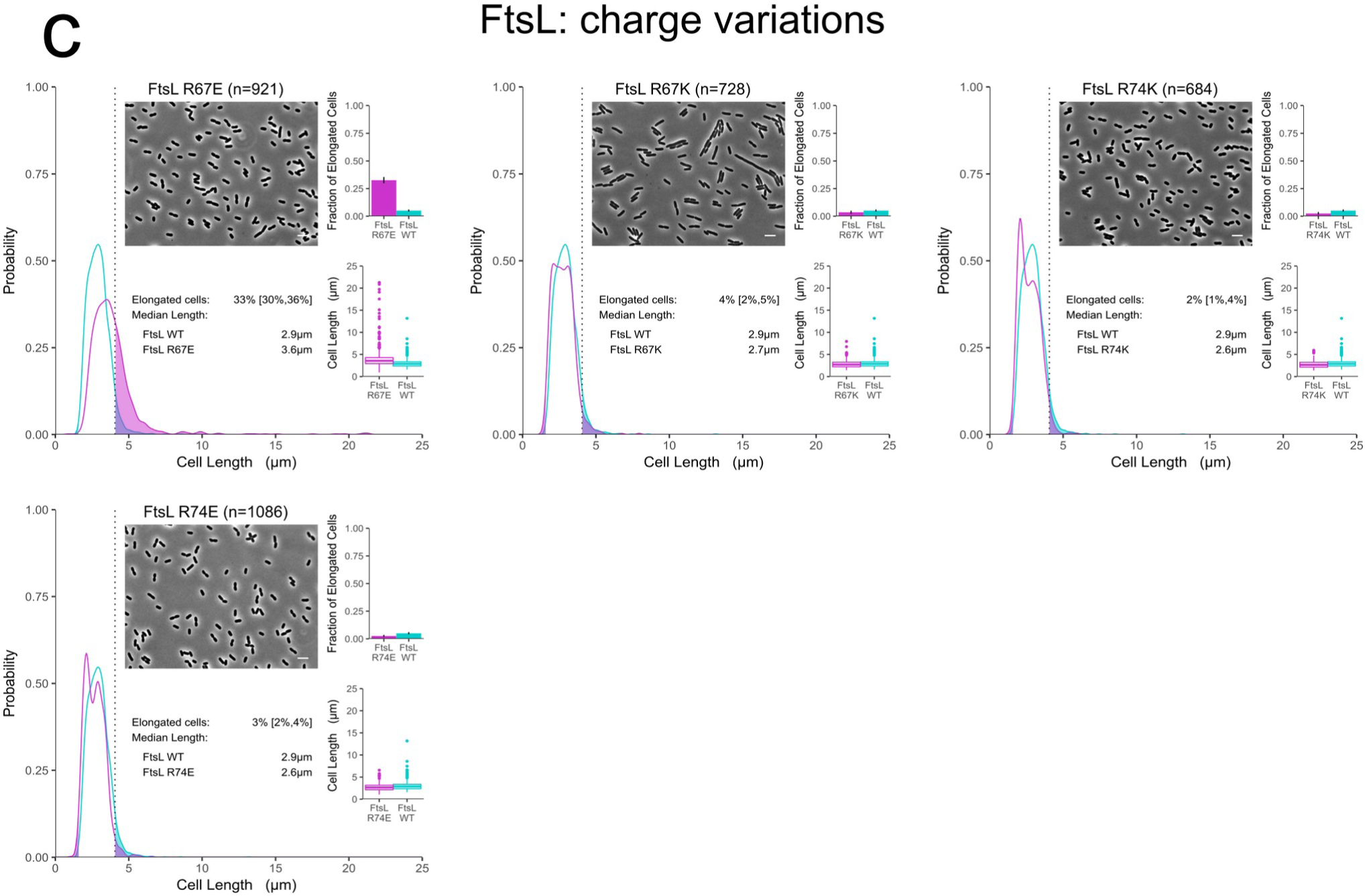
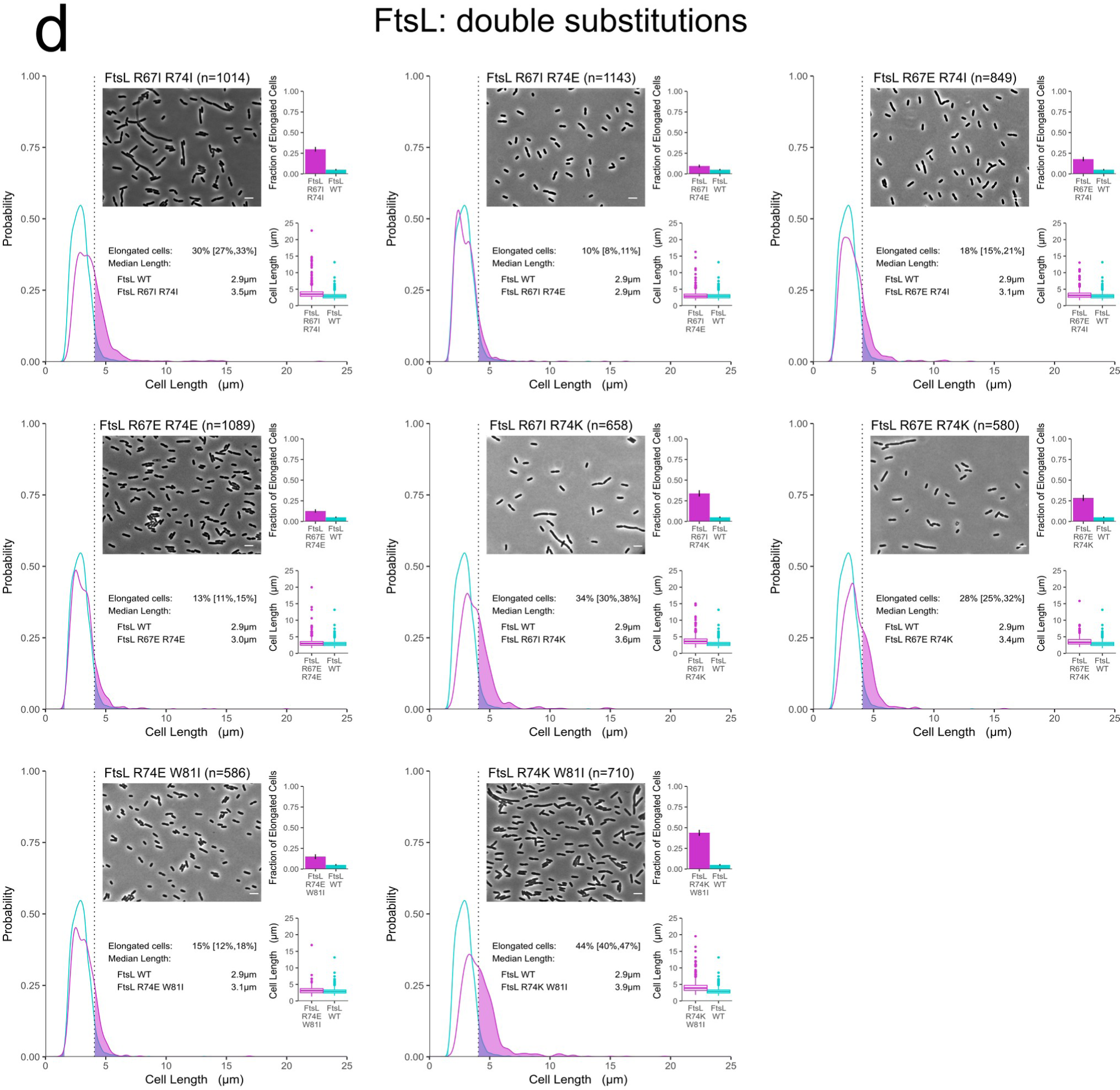
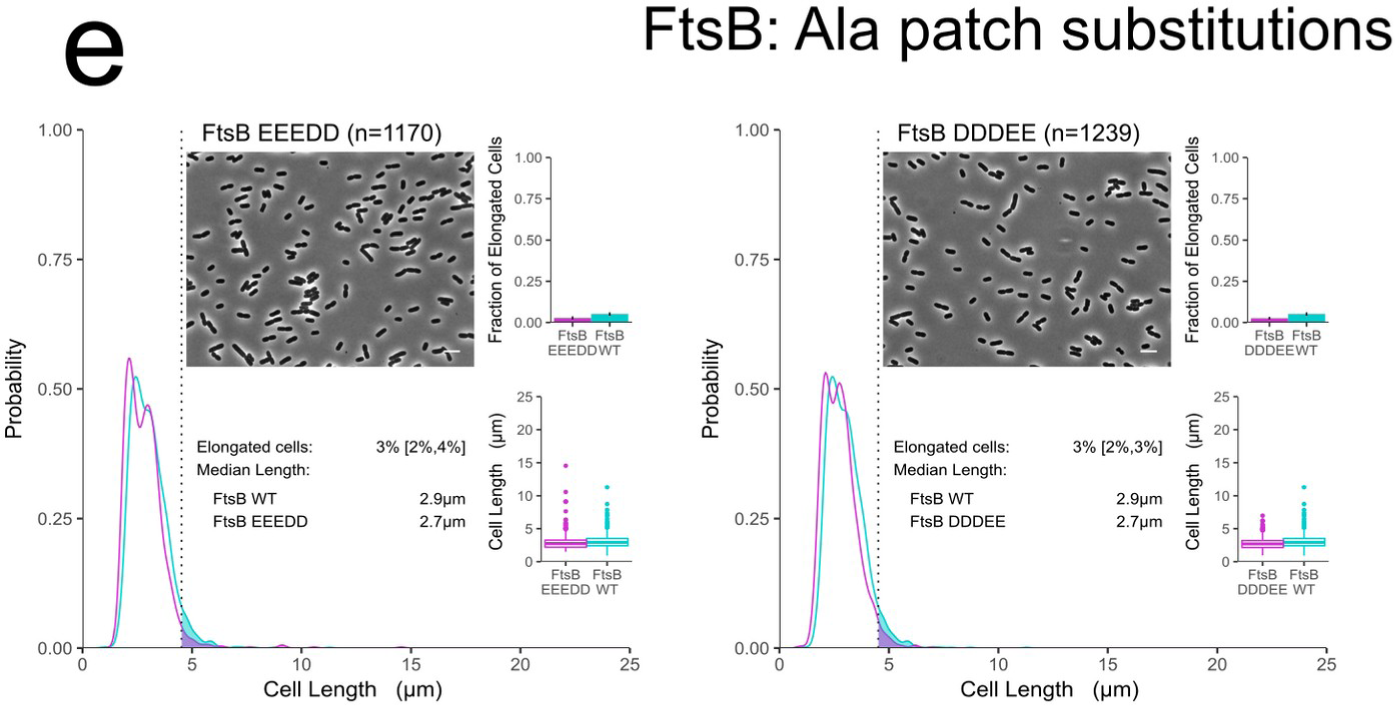
Cell length distribution of mutants. Phase-contrast images and cell length distributions of mutant cells (magenta) compared to wild-type (cyan). White scale bar = 5 μm. The shaded areas to the right of the dotted lines represent the fraction of cells that are longer than the 95th percentile in the WT distribution. The fraction is plotted as a histogram in the top inset. Error bars and bracketed values represent the 95% confidence interval for the fraction of elongated cells, estimated by 1000 bootstrap replications of the samples. The bottom inset is a box and whisker plot of the same distribution of the main panel. *a*) Nonideal to ideal mutations in FtsB. *b*) Nonideal to ideal mutations in FtsL. Next pages: *c*) Charge variations in FtsL at R67 and R74 positions. *d*) Double substitutions in FtsL at R67, R74, and W81 positions. *e*) Ala patch substitutions in FtsB in which all five Ala positions (37, 38, 41, 44, and 48) were simultaneously replaced with negatively charged residues, with a combination of three Glu and two Asp residues (EEEDD) and vice versa (DDDEE). Experiments were performed at 37 °C.

**Fig. S3.**
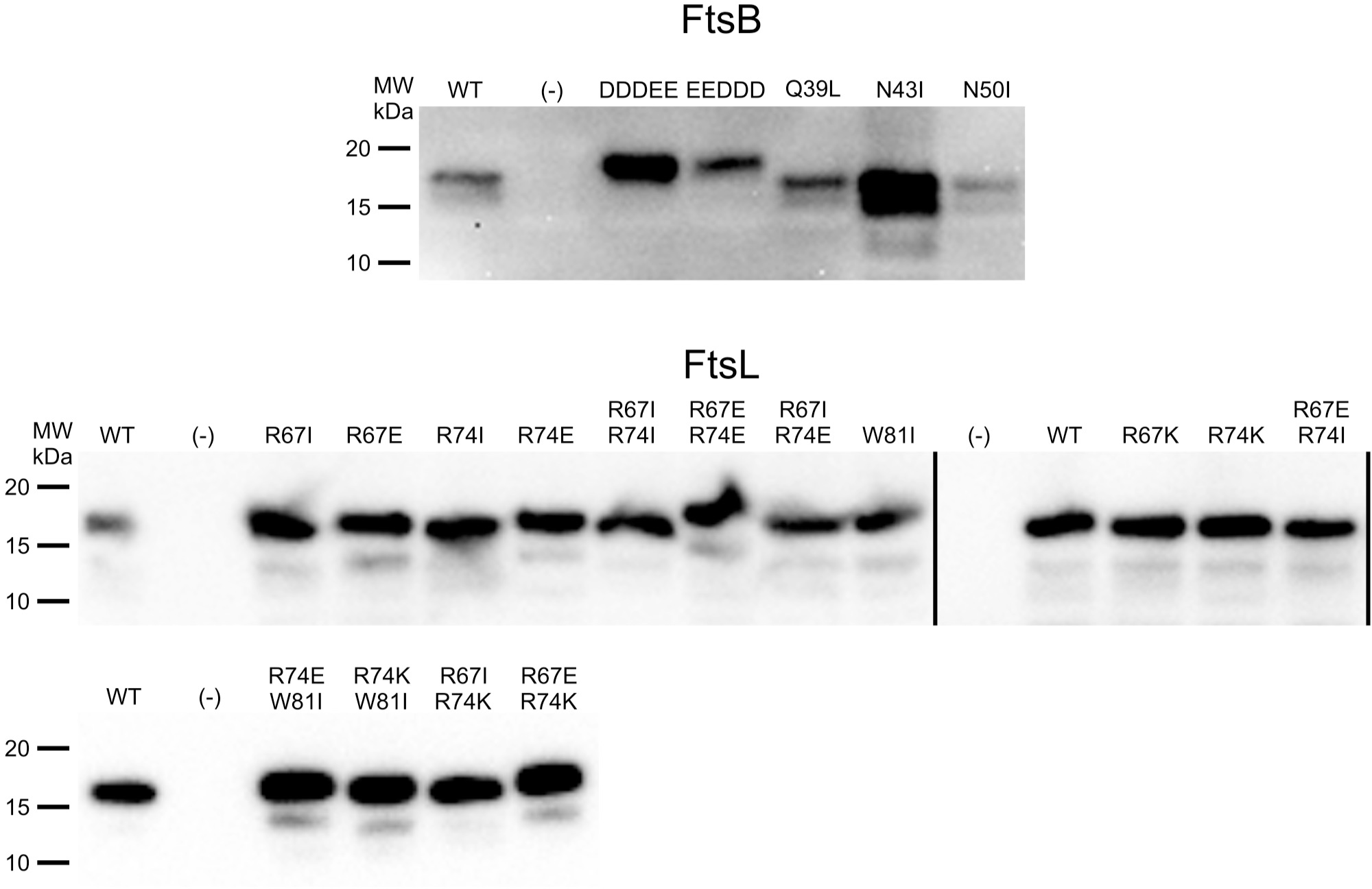
Expression level of FtsB and FtsL mutants assessed by Western Blot analysis. Image of all western blots of the FtsB and FtsL mutants tested in this work. Around twice as much whole cell lysate (normalized to total protein) was loaded for FtsB samples as for FtsL samples. Protein expression level of the FtsB and FtsL mutants with defective phenotypes are generally comparable to the respective wild type (WT). Negative controls (-) show no detectable signal for either protein. DDDEE is FtsB A37D/A38D/A41D/A44E/A48E, and EEEDD is FtsB A37E/A38E/A41E/A44D/A48D. Both show a slightly increased molecular weight, which may be due to the increased number of negatively charged residues in these mutants. There are cases of FtsB mutants with increased protein level (N43I, in particular), though it is unclear why. Individual gels are separated by solid lines.

**Fig. S4.**
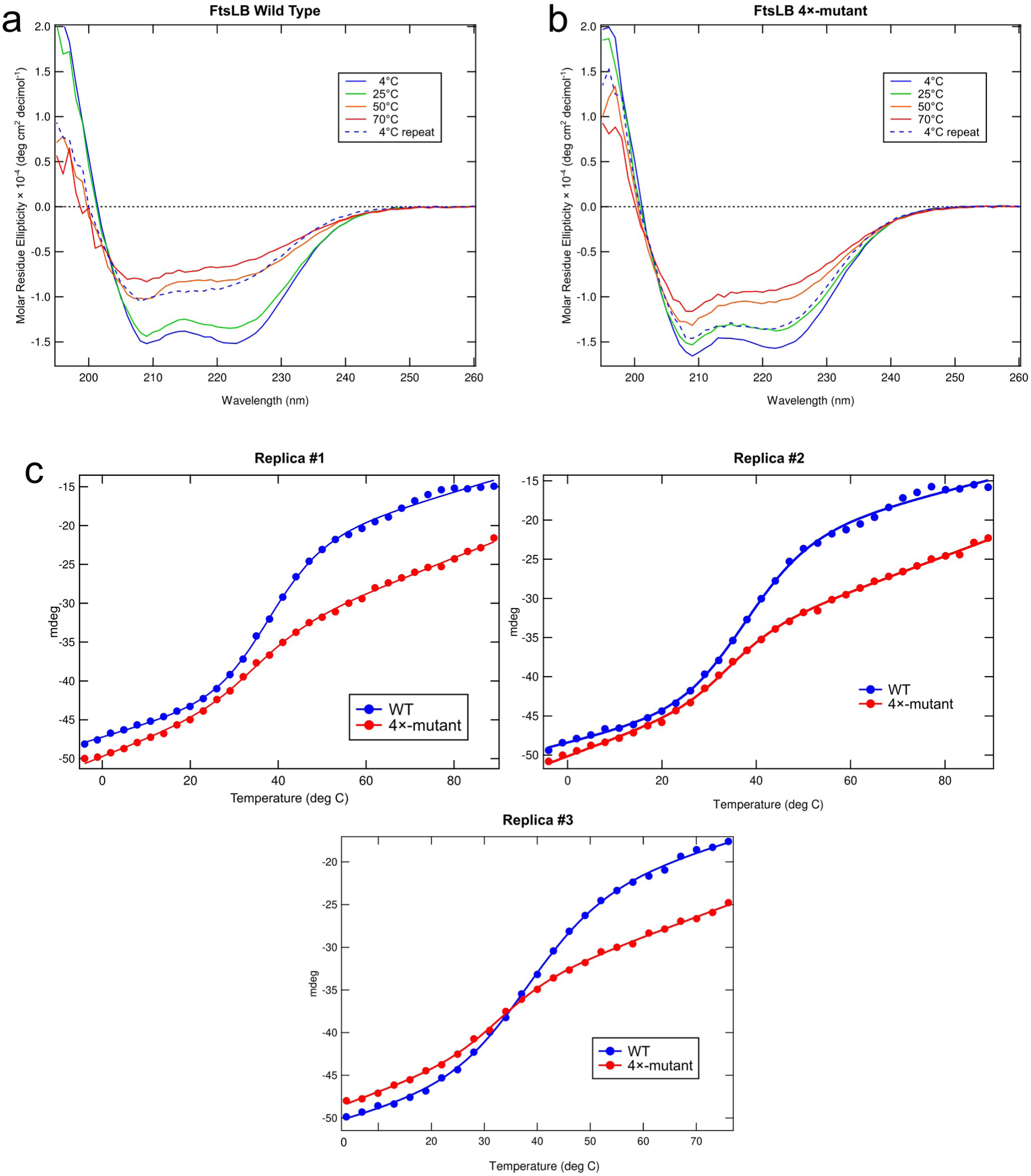
a) Far-UV CD spectra of WT FtsLB at 4, 25, 50, and 70 °C along with a 4 °C repeat after cooling the sample. b) Same analysis of the 4×-mutant. c) Replica experiments of temperature melting curves comparing WT FtsLB (blue) to the 4×-mutant (red). CD scans were monitored at 224 nm from 4°C to 89°C.

**Fig. S5.**
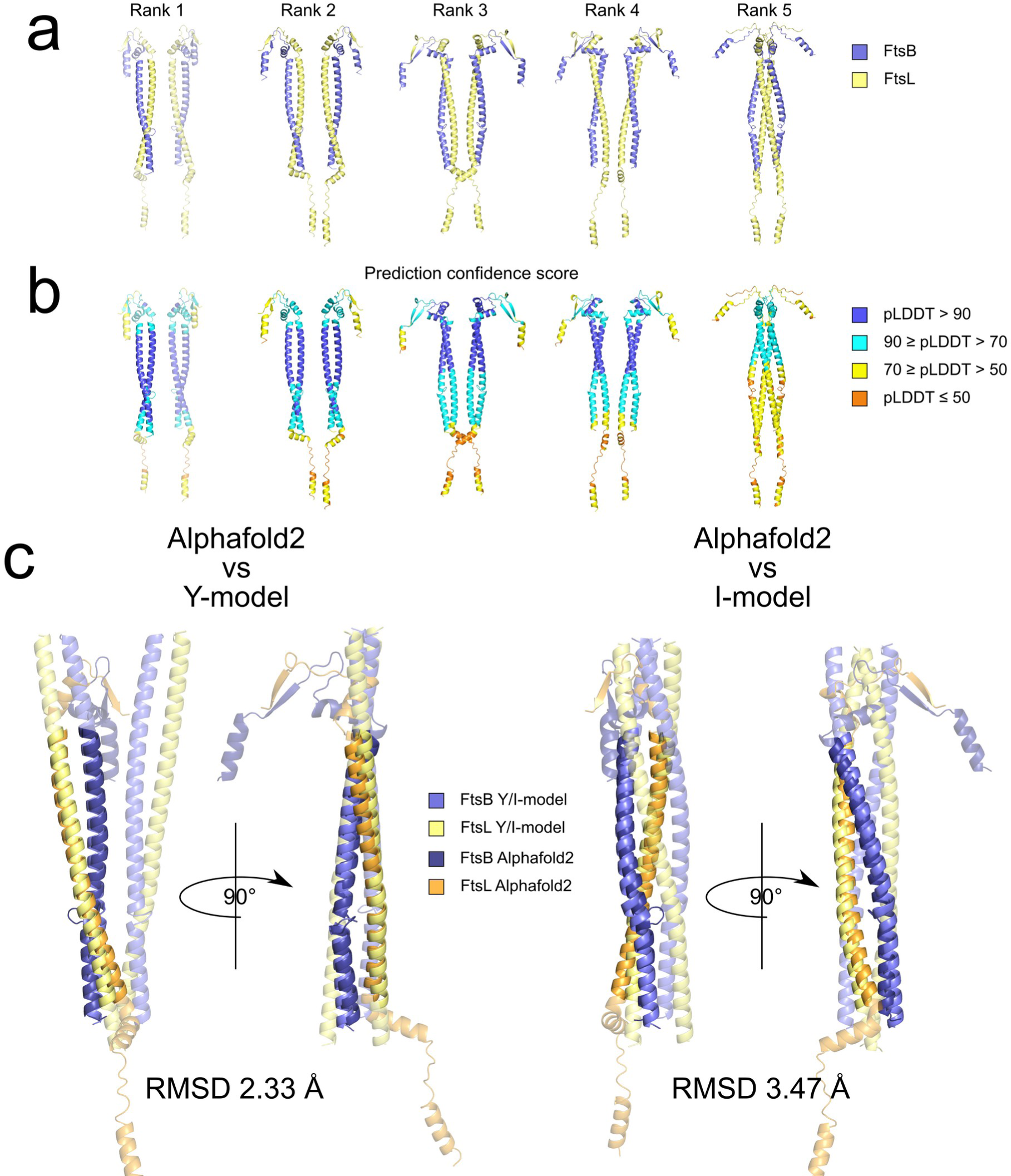
Alphafold2 models of FtsLB. a) Alphafold2 produced five ranked model of FtsLB, numbered 1-5 (best to worst). In spite of four subunits being provided, rank models 1-4 display separated dimeric units with little or no interaction between them. Rank model 5 is organized in a tetrameric unit. However, the interactions between the pair of dimers is loose and significantly underpacked. b) The Alphafold2 models have high confidence in the coiled coil region, as color coded in the figure (100: highest confidence; 0: lowest confidence). c) Alignments of one dimer from the rank 1 model to half of the Y-model (left) and the I-model (right). Regions not used in the alignment are transparent. The alignment with the Y-model is excellent, with a Cα RMSD of 2.33 Å, while the I-model alignment is less optimal, with an RMSD of 3.47 Å.

**Fig. S6.**
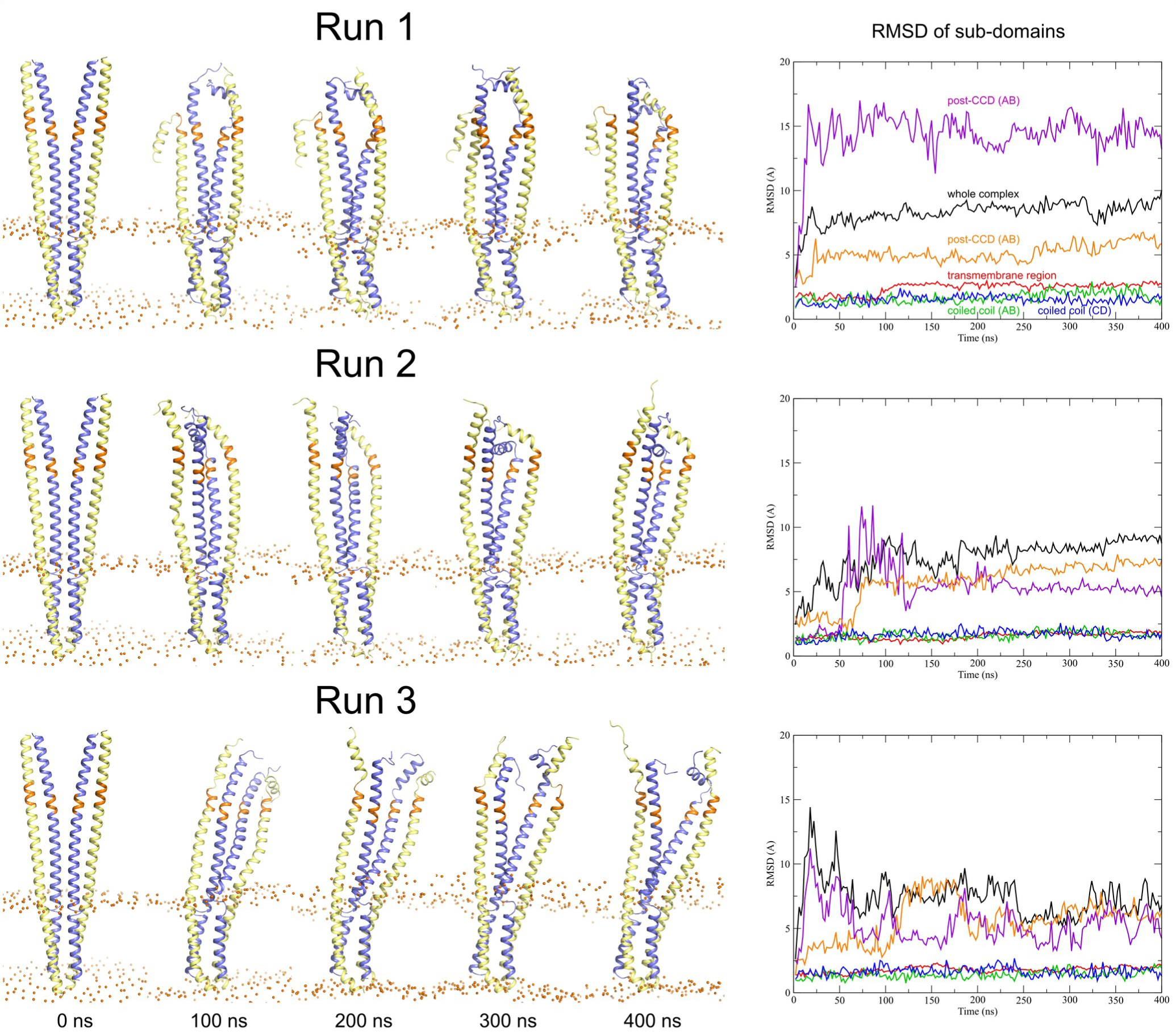
Trajectories of the three 400 ns replica MD runs of the FtsLB Y-model. The graph illustrates the fluctuations of the RMSD of the entire complex (black) and the individual subdomains: red: transmembrane region; green: coiled coil, chains A (FtsB) and B (FtsL); blue: coiled coil, chains C (FtsB) and D (FtsL); magenta: post-CCD region, chains A and B; orange: post-CCD region, chains C and D. The RMSD indicates that the transmembrane domain and coiled-coil domains remain relatively stable during the entirety of the simulations. The RMSD analysis is summarized in supplementary Table S3.

## Notes

### Competing Interest Statement

The authors have declared no competing interest.

### Summary of Updates

Minor revision of title and abstract

